# Ageing of the Upper Airway Epithelial Niche Limits Tissue-Resident T Cell Immunity

**DOI:** 10.64898/2026.07.10.737724

**Authors:** W. Huisman, L.A. King, Y. Singh, A.C. de Kroon, R.A.M. Steenbergen, I. van der Valk, R.S. Hagedoorn, E.J. de Meijer, M. König, T. Roek, G.E. Loe-Sack-Sioe, M.T. Wulffraat, M. van der Elst, S.L. Kloet, J.J.C. de Vries, A.H.E. Roukens, S.M. Kielbasa, C.E.W. Smeenk, D.O. Mook-Kanamori, P.S. Hiemstra, M. Yazdanbakhsh, M.H.M. Heemskerk, A.M. van der Does, S.P. Mooijaart, G.H. Groeneveld, S.P. Jochems

## Abstract

Respiratory tract infections are a major cause of morbidity and mortality among older adults worldwide. Yet, how ageing shapes protective immunity within the upper respiratory tract (URT) remains poorly understood. Here, we profiled the nasal immune landscape of young adults and older adults with and without frailty, using high-dimensional cytometry, proteomics, single-cell transcriptomics and T cell receptor (TCR) sequencing. Ageing was associated with a selective reduction of tissue-resident memory T (Trm) cells, independent of effector T cell numbers, inflammageing levels or frailty and was accompanied by a decline in T cell receptor repertoire stability. Trm cells from older adults also showed decreased steady-state IFNγ expression, accompanied by reduced antiviral transcriptional programs in the mucosa. Mechanistically, *in vitro* PBMC–epithelium co-culture models revealed that older adults have a reduced capacity for Trm differentiation, which was associated with a diminished epithelial TGF-β secretion, driven by reduced expression in ciliated epithelial cells. Together, these findings reveal that URT immunity deteriorates with age due to disrupted epithelial–immune crosstalk. This identifies the epithelial niche as a key regulator of mucosal immune ageing and suggests that enhancing TGFβ signaling may improve the effectiveness of mucosal airway vaccination strategies in older populations.

## Main

Respiratory viral infections represent a significant health burden among older adults, contributing to morbidity, hospitalization, and mortality worldwide^1, 2^. Age is one of the strongest risk factors for severe outcomes from common respiratory viruses such as influenza, respiratory syncytial virus and SARS-CoV-2^2–4^. In older adults, these infections can also lead to long-term complications, including a decline in functional capacity and the exacerbation of frailty, an age-associated syndrome characterized by diminished physiological reserves and increased vulnerability to stressors^3, 5^.

Ageing is accompanied by a gradual decline in immune competence, commonly referred to as immunosenescence. Importantly, accumulating evidence indicates that frailty is closely associated with features of immunosenescence^6^, suggesting that biological ageing of the immune system may be more advanced in older adults with frailty compared to their non-frail counterparts. One hallmark of immunosenescence is the reduced number and functionality of naïve T cells in the periphery, impairing the ability to mount robust responses to new pathogens and antigens^7^. This immunological decline has been linked to increased susceptibility to virus infections, increased disease severity and reduced vaccine responses in older individuals^8, 9^.

Emerging evidence highlights the pivotal role of tissue-resident memory T (Trm) cells in conferring protection to respiratory viruses and bacteria that enter via the upper respiratory tract (URT)^10–12^. Moreover, recent studies have shown that antigen-specific Trm cells can seed the URT mucosa in young adults after an infection^13–17^. Despite the well-documented systemic immune alterations with age, our understanding of how ageing impacts local immunity at mucosal surfaces remains limited.

In this study, we show that diminished epithelial TGF-β1 production in ciliated epithelial cells and reduced TGF-β1 sensitivity in T cells impair Trm differentiation in older adults, resulting in reduced URT Trm numbers and decreased clonal persistence across longitudinal timepoints. Moreover, decreased IFNγ-expression in Trm cells was accompanied by reduced antiviral gene signatures in the URT mucosa of older adults. Together, these findings reveal a key mechanism of age-related decline in mucosal immunity and reveal epithelial-immune crosstalk as a target in the design of airway mucosal vaccines to achieve durable protection in older populations.

## Results

### Age-related decline of tissue-resident memory cells in the upper-respiratory tract

To examine age-related changes in the mucosal immune system, we analysed participants from a prospective longitudinal cohort, including young adults (n=31) and older adults with (n=20) and without frailty (n=35). Older adults with frailty had significantly higher comorbidity scores, impairment of daily functioning and higher age compared with older adults without frailty **(Table 1 and Supplementary Table 1**). To assess which immune subsets from the URT were impacted by age and frailty, we performed phenotyping using a 34-marker spectral flow-cytometry panel of nasal immune cells that were obtained using minimally invasive nasal curettage. (**Figure 1a, Supplementary Table 2**). Unsupervised clustering on all immune cells (excluding granulocytes) resulted in 42 clusters, including 25 T cell, 2 B cell, 10 innate lymphoid and 5 myeloid clusters (**Figure 1b , Supplementary Fig. 1a and Fig. 2)**. None of the clusters were increased with age, while 10 cell clusters were significantly reduced, in both older adult groups. Nine of these decreased clusters were Trm clusters, and one was an atypical (CD11c^+^) tissue-resident memory B-cell (Brm) cluster. (**Figure 1c** and **Supplementary Fig. 1b-c**).

**Figure 1.**
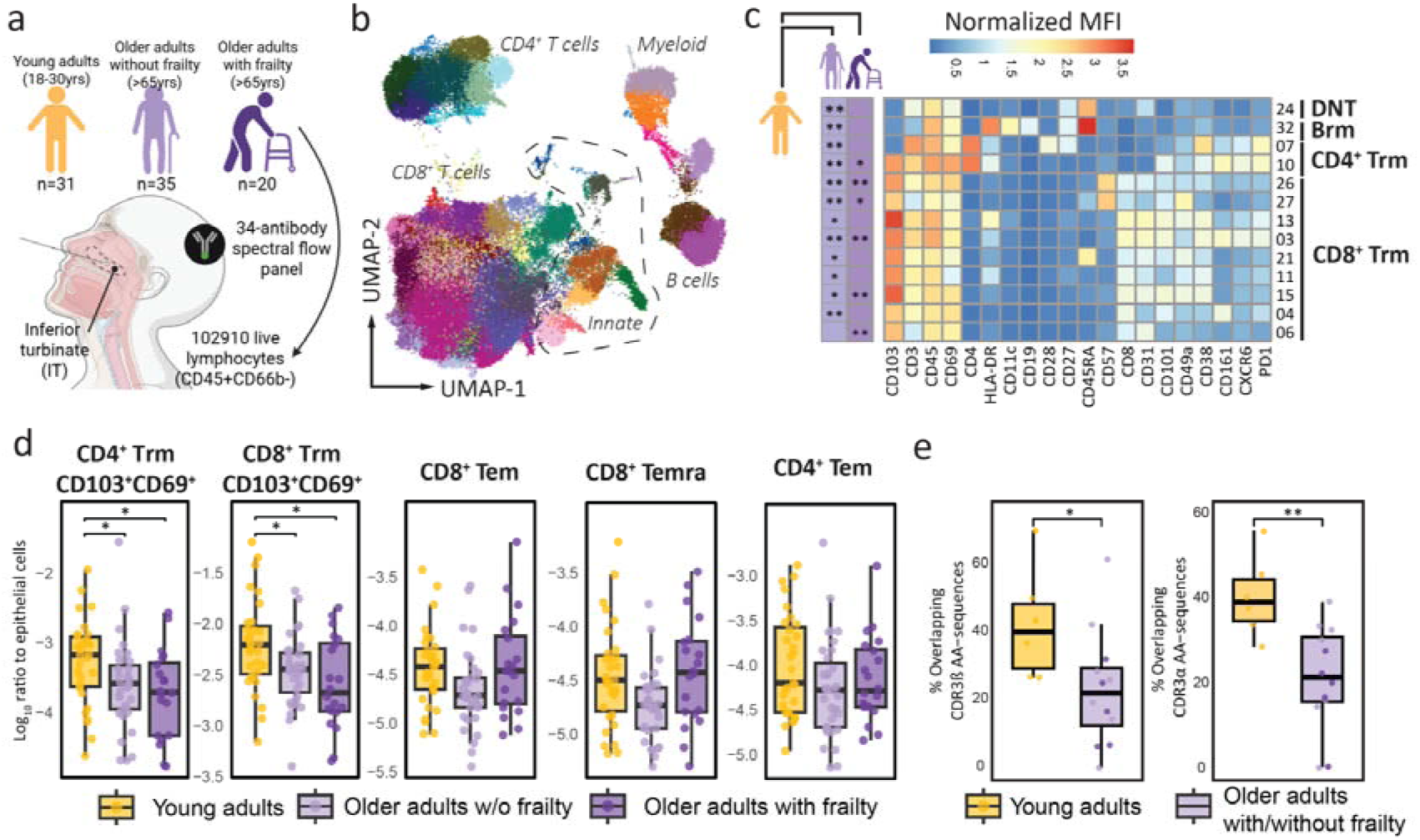
Age-related decline of immune cells in the upper-respiratory tract. Nasal-derived cells were collected using curettage and cryopreserved. **a**) Cross section of the upper respiratory tract, showing the anatomical site of sampling, the inferior turbinate. **b**) Uniform Manifold Approximation and Projection (UMAP) for all 102.910 nasal lymphocytes, excluding granulocytes. Unsupervised clustering was performed using FLOWsom resulting in 42 elbow clusters and was projected on the UMAP. **c**) Heatmap is shown of clusters (n=13) that were significantly different between groups. Only markers that were expressed by at least one of these clusters are shown. Statistical differences between young adults and older adults with/without frailty are displayed left of the heatmap and asterixis indicate level of significance. **d**) Manual gated T cell clusters, expressed as the log10 ratio of immune cells to epithelial cells. Data are presented as boxplots and compared between healthy young adults, older fit and frail adults. **e**) Bulk T Cell Receptor sequencing was performed to quantify clonal overlap and stability of T cells between paired samples collected at two timepoints 2-3 months apart with no self-reported intermittent respiratory tract infections. Overlap was assessed independently for CDR3α and CDR3β sequences, relative to the timepoint with fewest reads. Boxplots show the overlap for young adults and older adults with/without frailty with individual participants represented as jittered points. Statistical differences were assessed with Kruskal Wallis tests followed by Wilcoxon signed rank tests with Benjamini-Hochberg correction for multiple testing of multiple cell clusters and groups (C and D) *P < 0.05; ** P < 0.01, *** P < 0.001

**Figure 2.**
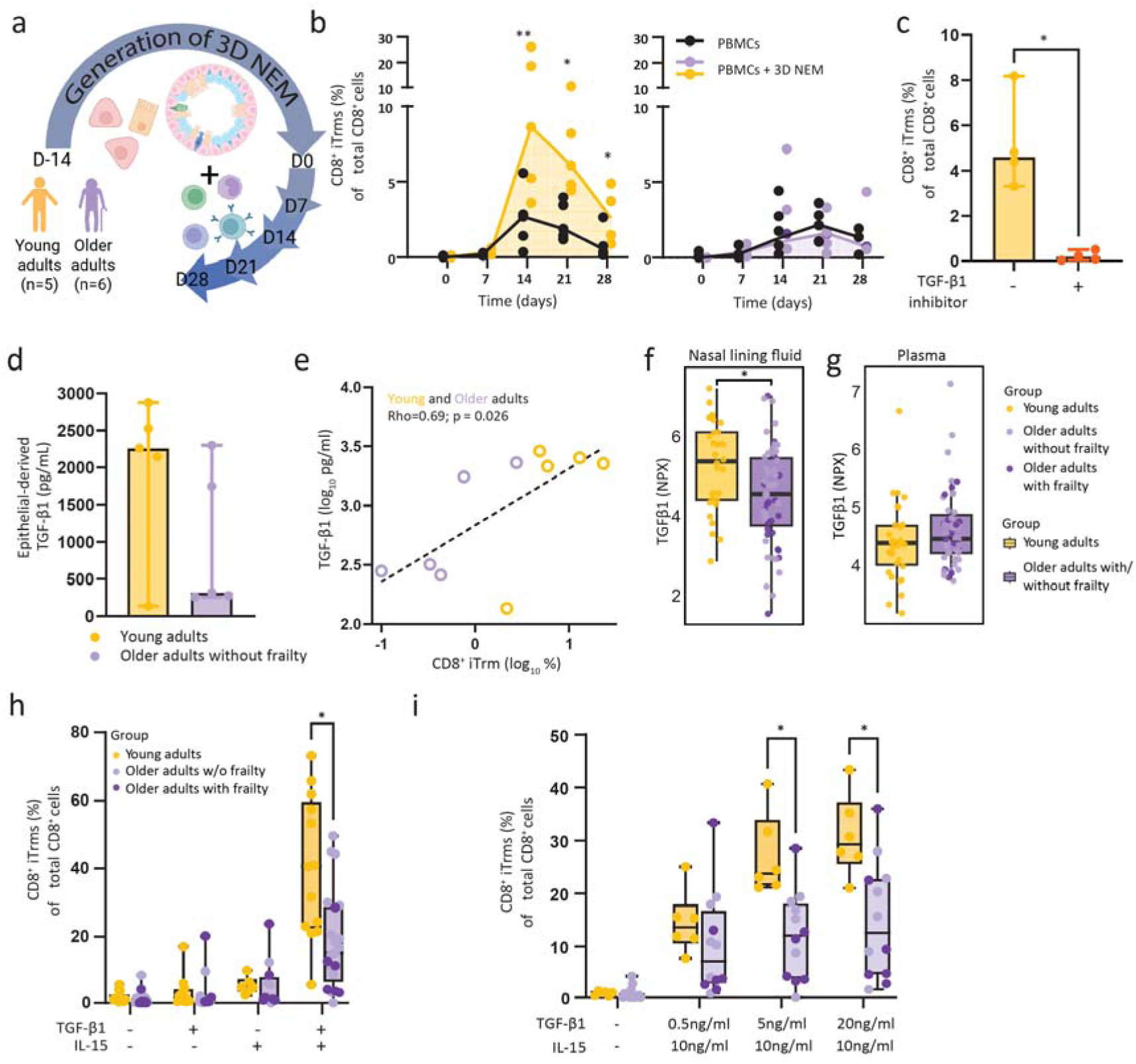
Impaired Tissue-resident memory T cell differentiation from older adults. The 3D nasal epithelial models (3D NEM) were generated from primary nasal epithelial cells collected from TINO participants using nasal curettage. After two weeks of differentiation, cells from 3D NEMs were harvested and co-cultured in the presence or absence of freshly thawed autologous PBMCs. **a**) Schematic overview of differentiation of epithelial cells to 3D NEM and co-culture with autologous PBMCs. At 7,14,21 and 28 days post co-culture initiation, induction of Trm phenotype was assessed by upregulation of CD103 and CD69. **b**) Percentages of CD8^+^ induced tissue-resident memory T cells (iTrm) over time are shown for PBMCs derived from young (n=5, yellow) and older adults (n=6, purple). PBMCs cultured without presence of 3D NEM are shown as black solid lines. Lines indicate medians. **c**) A 3D NEM was used where PBMCs were co-cultured from young individuals (n=4) and the percentage of iTrm is shown in the presence or absence of the TGF-β1 inhibitor SB431542 (10ug/mll) at day 21. **d**) Shown is the production of TGF-β1 (pg/ml) of epithelial cells from 3D NEMs from young (n=5) and older adults (n=5) at day 14. **e**) Spearman correlation plot is shown whereby the percentage of iTrm are shown against the epithelial 3D NEM production of TGF-β1. **f and g**) Relative abundance of TGF-β1 from nasal lining fluid (f) and plasma (g) using Olink. Protein levels are shown as NPX values (log_2_ scale). **h**) Barplots showing percentages of iTrm when PBMCs were cultured without 3D NEMs and supplemented with 5ng/ml TGF-β1 and/or 10ng/ml IL-15. **i**) Barplots showing percentages of iTrms when PBMCs were cultured without 3D NEMs and supplemented with IL-15 (10 ng/ml) and varying concentrations of TGF-β1 (0.5, 5.0, 20.0 ng/ml). Statistical differences were assessed with a linear mixed model (log_10_ iTRM% ∼ day × condition) (b), paired T test (c), Mann-Whitney U tests (d, h and i), or using two-group Wilcoxon rank-sum tests, with Benjamini–Hochberg correction applied across metrics to control the false discovery rate (f and g) *P < 0.05; **P < 0.01; ***P < 0.001; ****P < 0.0001

**Table 1.** Cohort characteristics. For continuous variables, median and interquartile range (IQR) are presented, and a two-sided Kruskal–Wallis test with Benjamini–Hochberg correction for multiple comparisons was used for statistical comparisons of 3 groups whereas a Mann-Whitney U test was performed when older fit adults were compared to older adults with frailty. ^#^this group was excluded for statistical analysis. *P < 0.05, **P < 0.01, ***P < 0.001, otherwise nonsignificant. ^a^Dunn post hoc test relative to young adults, P < 0.05. ^b^Dunn post hoc test relative to healthy older adults, P < 0.05. For categorical variables, number of participants and percentages are listed in the subgroup are presented, and a χ2 test was used for comparison. BMI: body mass index; NA: not applicable

|  | Young Adults | Healthy older adults | Frail older adults |
| --- | --- | --- | --- |
| **Patients included** | 31 | 35 | 20 |
| **Sociodemographic** |  |  |  |
| Age (years old) ** | 27 (24-29)<sup>#</sup> | 77 (73-79) | 81.5 (77-89) |
| Sex, n male (%) | 14 (45%) | 17 (49%) | 8 (40%) |
| **Anthropometrics** |  |  |  |
| Weight (kg) | 73 (65-84) | 69 (63-80) | 69 (62-94) |
| Height (cm) ** | 180 (170-185) | 170 (164-176)<sup>a</sup> | 162.5 (158-180)<sup>a</sup> |
| BMI (kg/m<sup>2</sup>) * | 23.31 (21-25) | 24.49 (22-26) | 26.07 (23-28)<sup>a</sup> |
| Handgrip strength (kg) | NA | 28 (22-38) | 27 (16-31) |
| Men | NA | 38 (33-41) | 33 (26-40) |
| Women | NA | 23 (18-27) | 20 (14-27) |
| **Medical data** |  |  |  |
| Co-morbidity score*** | NA | 3 (3-5) | 5 (4-6) |
| Clinical Frailty Score*** | NA | 1 (1-2) | 5 (4-6) |
| KATZ-ADL score*** | NA | 0 (0-0) | 1 (0-1) |
| KATZ-iADL score*** | NA | 0 (0-3) | 5 (2-7) |
| 6-Cognitive Impairment Test (6-CIT) | NA | 2 (0-4) | 2 (0-5) |
| Past smoking (<40 packyears), n (%) | 5 (16%)<sup>#</sup> | 15 (43%) | 9 (45%) |

The CD8^+^ Trm subclusters that were significantly decreased with age all expressed CD31, associated with trans-endothelial migration^18^, and/or CD57, a marker of antigen-driven differentiation^19^. Manual gating confirmed a decrease of the entire tissue-resident CD103^+^CD69^+^ T cell population for both CD4^+^ and CD8^+^ T cells, while no differences were observed for any of the effector memory T cell (Tem) populations (**Figure 1d, Supplementary Fig. 3 and 4a**). To assess whether this Trm decline was linked to inflammageing, we quantified systemic circulating cytokines associated with age-related chronic inflammation, including interleukin-6 (IL-6). Inflammageing was evident in fit and frail older adults and plasma IL-6 levels were also significantly associated with frailty after adjusting for age and sex (Log_2_-Est=0.99, *P=0.014*; **Supplementary Fig. 4b,c**).

However, the reduction in Trm cell numbers did not correlate with any inflammageing-related cytokines, nor was it associated with frailty (**Figure Supplementary Fig. 4d-f).**

We hypothesized that the reduced Trm numbers in ageing individuals could result from impaired recruitment of Tem cells to the URT in response to prior infections, impaired differentiation of Tem cells into Trm cells and/or reduced maintenance of already differentiated Trm. To assess whether older adults exhibit impaired recruitment of Tem, CyTOF was performed on nasal curettage samples from 65 individuals with acute SARS-CoV-2 infection (median age: 54 years, IQR: 34–70; median time post-infection: 11 days, IQR: 8–15) (**Supplementary Fig. 5; Supplementary Table 3**). Effector cell populations were increased during acute SARS-CoV-2 infection in older adults, suggesting that recruitment of circulating T cells to the mucosa is preserved (**Supplementary Fig 5 and 6**). As Trm cells were reduced in both older adults with and without frailty, and effector cell recruitment was intact, we assessed temporal stability of URT T cells by longitudinal T Cell Receptor (TCR) sequencing of nasal curettage samples from 18 individuals 2-3 months apart. In young adults, a median of 40% of TCRβ sequences were shared between the two timepoints, indicating relatively stable T cell populations. In contrast, this overlap dropped to 22% in older adults (*P=0.024*), indicative of a reduced T cell repertoire stability with age. A similar decline was observed for TCRα sequences (*P=0.0065*; **Figure 1e**). TCR repertoire evenness was comparable between groups, as reflected by similar Shannon diversity and Inverse Simpson indices (**Supplementary Fig. 7)**.

These data indicate that ageing associates with reduced Trm numbers and a concomitant reduced stability of the T cell repertoire in the URT mucosa, while effector cell recruitment remains intact. However, Trm reduction does not associate with other canonical ageing processes such as inflammageing and frailty.

### T cells from older adults exhibit impaired Trm differentiation in a 3D nasal epithelial model

To test whether Trm differentiation is impaired in older adults, we established a human-derived three-dimensional nasal epithelial model (3D NEM) that supports long-term co-culture with autologous PBMCs^21^ (**Supplementary Fig.8a**). This co-culture leads to the *in vitro* induction of Trm-like cells, (iTrm), which phenotypically, transcriptionally and functionally resemble Trm cells^21^. Using this model, we evaluated the capacity of peripheral blood-derived T cells from young (n=5) and older adults (n=6, 5 without and 1 with frailty) to differentiate into iTrm cells (**Figure 2a**). In the presence of epithelial cells, PBMCs from young adults differentiated into CD69^+^CD103^+^ CD8^+^ iTrm cells, peaking at day 14 (median 8.6%, *P = 0.0025* relative to PBMCs only) and remained elevated at days 21 and 28 (**Figure 2b and Supplementary Fig. 8b)**. In contrast, PBMCs from older adults showed no significant differentiation into iTrm (median 1.3%, *P*=0.40), comparable to cultures without epithelial cells.

Similarly, CD4^+^ iTrm were induced in young adults, but not older adults, in the presence of epithelial cells (**Supplementary Fig. 8c**). Given that TGF-β1 is reported to be an important driver of the differentiation of Tem into Trm in most tissues^22^, albeit not in the URT mucosa in a murine model^10^, we blocked the TGF-β1 receptor in our co-culture model. This completely prevented iTrm differentiation in young adults, demonstrating that TGF-β1 is critical in this human co-culture system (**Figure 2c**). TGF-β1 was produced at lower levels by epithelial cells derived from older adults compared with those derived from young adults (median = 319 pg/mL versus 2261 pg/mL, *P=0.31*, **Figure 2d**). Moreover, TGF-β1 production by epithelial cells from the 3D NEM positively correlated with the frequency of CD8[ iTrm cells (rho=0.69, *P*=0.026, **Figure 2e**). No differences were observed in IL-15 production, a survival factor for Trm^23^ , between epithelial cells of young and older adults (**Supplementary Fig. 8d**). In line with these *in vitro* results, *ex vivo* analysis of nasal lining fluid showed significantly reduced levels of TGF-β1 in older adults compared to young adults (**Figure 2f** and **Supplementary Fig. 9 and 10**). In contrast, plasma TGF-β1 levels were not decreased in older adults, showing compartmentalization of the effects of ageing and confirming the critical role for local TGF-β1 production (**Figure 2g**).

To determine whether impaired iTrm generation in older individuals could be rescued by exogenous TGF-β1, we supplemented PBMC cultures from young, older adults with and without frailty with TGF-β1 and/or IL-15. PBMCs from older adults with and without frailty exhibited iTrm differentiation in response to TGF-β1 and IL-15, but this was significantly lower in older adults than young adults (**Figure 2h and Supplementary Fig. 8e**), indicating also a T cell intrinsic impairment in the capacity to generate iTrm. The response to TGF-β1 was dose-dependent in young adults but reached a plateau early in older adults (**Figure 2i)**. The reduced T cell sensitivity of older adults to TGF-β1 could potentially be explained by lower levels of the TGF-β-receptor on T cells. In a publicly available single-cell dataset^24^, the expression of TGF-β-receptor II in naïve and central memory T cells was indeed decreased in older adults (n=45; 55-65 years) compared to young adults (n=49; 25-35 years) (**Supplementary Fig.8f**).

In conclusion, T cells from older adults show reduced responsiveness to TGF-β1, which, combined with lower TGF-β1 production from epithelial cells from older adults, prevents Trm differentiation.

### Single cell RNA sequencing reveals age-associated transcriptional changes in the URT

To understand how ageing reshapes the transcriptional landscape of nasal Trm cells and to pinpoint which epithelial subsets contribute to TGF-β1 production in the mucosal niche, we applied single-cell RNA sequencing to nasal curettage samples of healthy young and older adults with and without frailty (n=30, 10 per group; **Supplementary Table 7**). In total, 9 different immune and 10 different epithelial cell-populations were defined using clustering (**Figure 3a and 3b**).

**Figure 3.**
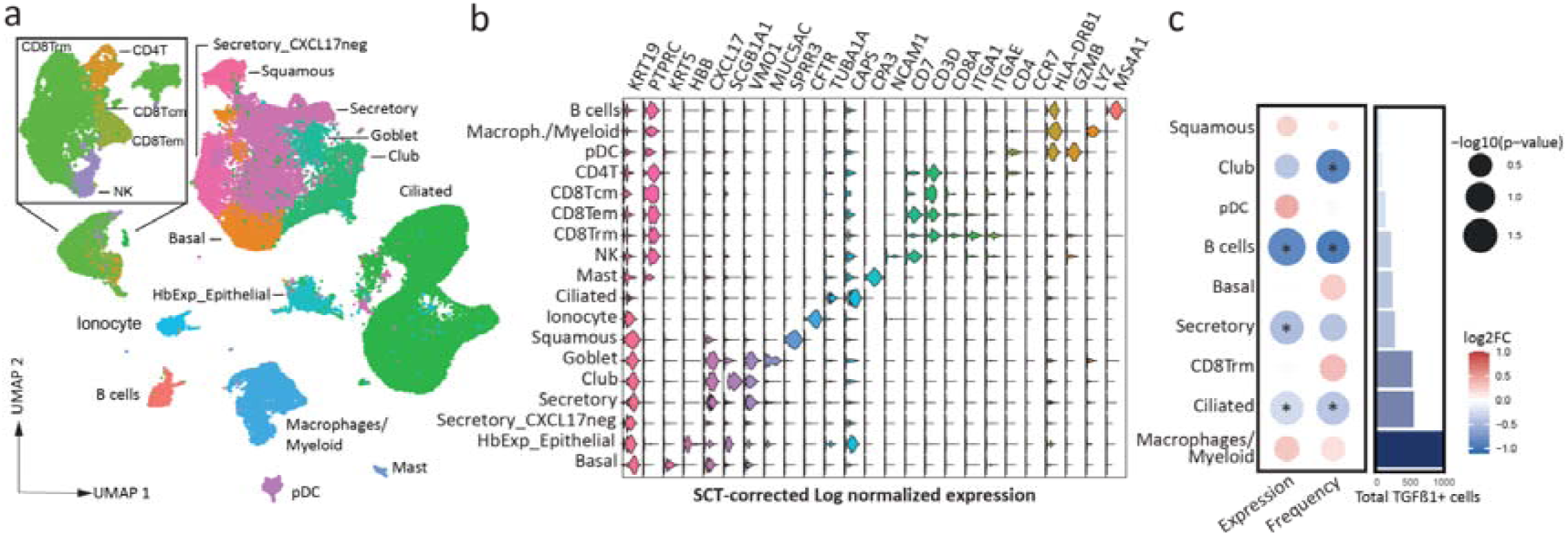
Single cell RNA sequencing reveals age-associated transcriptional changes in TGFB1 expression of ciliated epithelial cells. **a**) UMAP visualization of all 19 immune and epithelial populations identified by single-cell RNA-sequencing of nasal curettage samples (n=30, 66.991 cells). **b**) Violin plots showing population-specific marker gene expression. **c**) Bubble heatmap showing log_2_ fold change of TGFB1 expression and TGFB1L cell frequency between older adults and young adults across cell types. Cell frequencies refer to the proportion within each cell population. Only nasal cell populations with at least 100 TGFB1 reads were assessed. Bubble color encodes foldchange direction, with blue indicating lower values in older adults and red indicating higher values in older adults. Horizontal bar plots depict the number of cells expressing TGFB1 per population.

**Figure 4.**
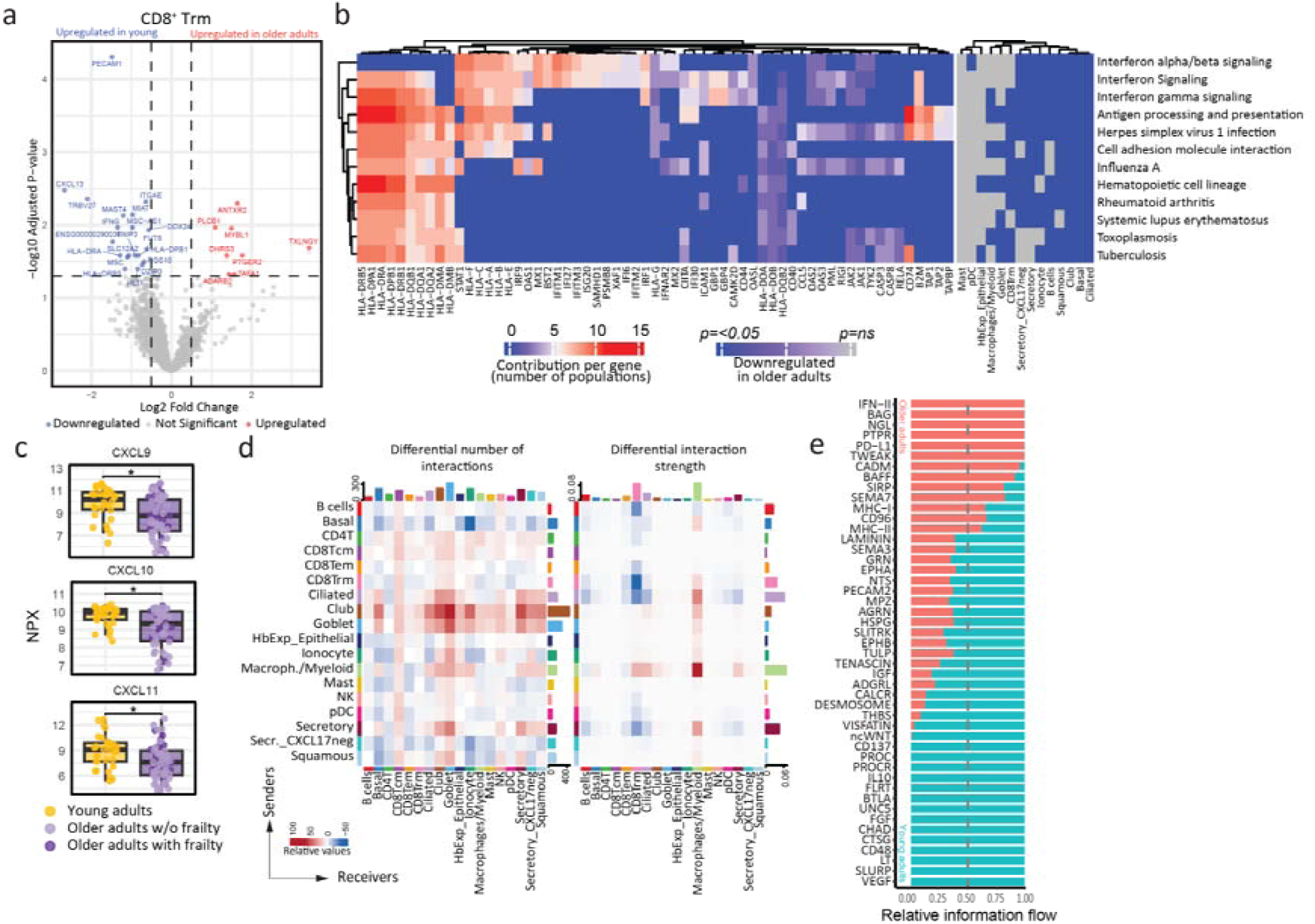
**Reduced CD8 Trm IFN**γ **anti-viral programming in the URT of older adults. a**) Volcano plot of differentially expressed genes (DEGs) in CD8L Trm cells, highlighting transcriptional pathways altered with ageing based on pseudobulk analysis using limma-voom. Blue indicates lower expression in older fit and frail adults, whereas red indicates higher expression in older fit and frail adults). **b**) Significantly enriched GSEA pathways that are downregulated in older fit and frail adults across different cell populations are shown, together with their leading-edge genes. Leading-edge genes represent the core subset of genes that drive the enrichment of each pathway, contributing most strongly to the observed downregulation in older fit and frail adults. **c**) Relative abundance of IFN-g induced proteins CXCL9-11 from nasal lining fluid using Olink. Protein levels are shown as NPX values (log_2_ scale). **d**) Differential cell-cell communication between young and older fit/frail adults visualized by CellChat. Heatmaps display global differences in inferred ligand–receptor interaction strength across all annotated cell populations. The left and right panel show the differential interaction probabilities and strengths respectively, where red indicates increased interactions in samples from older fit/frail adults and blue indicates increased interactions in young adults. **e**) Stacked barplots summarizing the relative contribution of each signaling pathway to the overall communication network. Significance testing was performed within CellChat and pathways with adjusted p-value < 0.05 and at least 50% difference between groups are shown .

Notably, frequencies of immune cell populations identified by single-cell RNA sequencing showed good concordance with abundances measured by spectral flow cytometry (**Supplementary Fig. 11a-b and Fig. 12**). Single-cell sequencing further enabled the assessment of age-associated changes in epithelial cell subsets, revealing an increased abundance of club cells in older adults compared to young adults, while ionocytes were specifically enriched in adults with frailty (**Supplementary Fig. 11a and Fig. 12**). Ciliated epithelial cells were the predominant source of *TGFB1* from epithelial cells in the nasal mucosa, together with macrophages from the immune compartment (**Figure 3c**). A statistically significant reduction in proportion of *TGFB1*[ cells was observed specifically in ciliated epithelial cells, club cells, and B cells in older adults, with no changes detected in any of the other subsets. (**Figure 3c and Supplementary Fig. 13a and 14**). Pseudobulk differential expression analysis confirmed lower *TGFB1* expression by ciliated epithelial and B cells, with secretory epithelial cells additionally showing significant reduction in expression of *TGFB1* (*P*=0.041) in older adults. The TGF[β1 protein levels measured in nasal lining fluid positively correlated with *TGFB1* expression levels for ciliated cells, but not for macrophages, B cells, or secretory cells (**Supplementary Fig. 13b)**.

Taken together, ageing was associated with alterations in epithelial cell composition, including reduced *TGFB1* expression in ciliated epithelial cells, a major source of TGF-β1 within the URT mucosa.

### Reduced CD8 Trm IFN**γ** anti-viral programming in the URT of older adults

Differential gene expression analysis of nasal cell populations between young and older adults was performed to identify further age-related transcriptional changes, revealing substantial changes in several populations, including CD8[ Trm cells (**Figure 4a**).

Consistent with our flow-cytometry results, expression of *PECAM1* (CD31) and *ITGAE* (CD103) were significantly reduced in older adults, as was *IFNG*. Secreted IFNγ levels were also lower in nasal lining fluid of older adults (**Supplementary Fig.10**). Gene set enrichment analysis was then performed for each of the nasal populations to identify age-associated pathways. Interferon signaling, cell-adhesion molecule interaction, antigen processing and pathways associated with viral infections such as Influenza A were consistently downregulated in older adults across nasal cell populations (**Figure 4b**). In line with this broad downregulation of IFNy-induced pathways, protein levels of typical IFNy-induced antiviral cytokines CXCL9-11 were also shown be decreased in nasal lining fluid of older adults (**Figure 4c**). Cell-cell communication using ligand-receptor analysis with CellChat revealed a marked age-associated rewiring of epithelial-immune interactions.

In young adults, CD8[ Trm cells engaged strongly with other immune and epithelial cell subsets (**Figure 4d**). One of the important drivers for this was IFNy (IFN-II) signaling, which was dominated by CD8^+^ Trm and diminished with age (**Figure 4e, Supplementary Fig. 15 and 16**).

Older adults instead showed enhanced myeloid cell signaling towards Trm and ciliated epithelial cells.

Together, these data reveal an age-associated disruption of epithelial–Trm homeostatic interactions, characterized by altered epithelial TGF-β1 dynamics and diminished CD8^+^ Trm IFNγ-driven antiviral gene expression pathways in older adults.

## Discussion

Here, we found that the number of Trm cells in the URT mucosa are specifically decreased in older adults. Mechanistically, recruitment of effector cells during acute infection was intact, but a reduced sensitivity to TGF-β coupled with decreased TGF-β production by predominantly ciliated epithelial cells were associated with an impaired capacity to differentiate into Trm cells. This reduction in Trm is consistent with the lower stability of URT mucosa T cell clones, indicating a reduced capacity to maintain tissue-resident immunity in older adults. These findings suggest that the ageing epithelial niche may contribute to mucosal immune decline, highlighting a potential role for tissue environments in constraining tissue-resident immunity.

While recent studies have shown that antigen-specific T cells can seed the URT mucosa, the longevity of this tissue-resident immunity remains an open question. In mice, nasal Trm can persist for months^10^, which was supported by a recent study in humans that showed that influenza-specific T cells can persist in the nasal cavity for at least six months^13^. In addition, very few studies have assessed the effect of age on mucosal immunity. Nguyen et al showed in a pilot study of 8 individuals that the frequency of CD8^+^ Trm, but not other cell types in the lung decreases with age^25^. Urban et al showed that total nasal CD8^+^ and CD4^+^ T cells were decreased in older adults^26^, although the underlying mechanisms and subsets affected were not analyzed. Here we observed that in young adults 40% of TCRβ chains were still present in the upper respiratory tract after 2-3 months, quantifying the stability of the nasal T cell pool. However, with age, this overlap decreased to 22%. This decreased TCR stability in the nasal mucosa of older adults contrasts with earlier findings in blood, where the repertoire stability was found to be higher in older adults than young adults^27^, highlighting the relevance of studying local immune responses.

The reduction of Trm cells observed in older adults was not related to either frailty or to systemic markers of inflammaging, suggesting an alternative axis of immunological aging that thus far has been underappreciated. The underlying impaired TGF-β production was not visible in blood, but derived from changes in the epithelial niche. In addition, we observed changes in club cells and ionocytes related to aging and/or frailty, highlighting the need for further studies into how epithelial cells are affected by ageing^32^ and the functional consequences of tissue-ageing.

The reduced ability to generate Trm cells has consequences for the induction of local immunity after infection or nasal vaccination. The live attenuated influenza vaccine (LAIV) shows reduced efficacy in older adults, and is therefore only licensed up to 49 years of age in the US. The current dogma is that increasing baseline immunity with age blocks replication of the vaccine, and in this way impairs LAIV immunogenicity^28^. However, as this vaccine depends on the induction of tissue-resident memory^29^, our findings suggest that an inability to generate local immunity could contribute to reduced efficacy. More broadly, this suggests that any URT mucosal vaccination strategy would need to incorporate strategies to overcome the reduced Trm differentiation to achieve protection in older adults.

The Trm that were generated in older adults produced lower levels of IFNγ in homeostasis compared to young adults. IFNγ has been shown to be central for the antiviral effects of Trm in a mouse model of Sendai virus infection^11^. Cell-cell signaling and pathway analysis indicated that this IFNγ reduction in older adults led to reduced baseline levels of anti-viral responses and lower antigen-presentation capacity of nasal immune and epithelial cells. This finding offers a mechanistic explanation for large scale studies demonstrating that SARS-CoV-2 viral loads are higher in older adults^30^. It is also in line with mathematical modeling of daily SARS-CoV-2 kinetics, which has suggested that with age the ability to induce refractory cells via interferons decreases^31^.

This study has limitations. we were unable to longitudinally follow sufficient young and older adults post infection to track antigen-specific T cell clones, and our Trm stability data is therefore based on the total TCR repertoire. Moreover, as epithelial-immune co-culture requires an autologous setting, we were unable to perform experiments where PBMCs from younger adults were combined with epithelial cells from older adults and vice versa. Finally, as we focused on discrete groups of young and older adults, we cannot pinpoint an age-range in which Trm formation starts to be impaired.

In conclusion, using a combination of *in vitro* co-culture models, bulk TCR-sequencing, high-dimensional flow and mass cytometry, proteomics and single-cell transcriptomics, we revealed a mechanism where Trm differentiation is impaired with age due to reduced TGF-β signaling, with alterations in the epithelial compartment restricting tissue-resident immunity in the URT in older adults. This suggests a possible impact on susceptibility to infection, and pinpoints targets that can be used to improve airway mucosal vaccination strategies in older adults.

## Methods

### Study participants

All participants provided written informed consent according to the Declaration of Helsinki. Samples obtained from the URT by nasal curettage were obtained as part of the TINO clinical trial (https://clinicaltrials.gov/study/NCT06039527), a prospective cohort study investigating the mechanisms and underlying cause of T cell decline in the upper-respiratory tract that was conducted at the Leiden University Medical Center (LUMC, Netherlands). The study included young healthy individuals aged 18-30 years, older adults with and without frailty (>65 years). Frailty status was determined using the Clinical Frailty Scale (CFS) with a score of 4 or higher indicating frailty^33^. The young individuals were recruited in the western part of the Netherlands, the older individuals were recruited in two general practices in the western part of the Netherlands. In and exclusion criteria can be found in **Supplementary table 1**. In summary, participants were excluded in case of inability to provide informed consent or with any type of chronic respiratory-related illness such as asthma, COPD or chronic rhinosinusitis. Participants with a recent history of vaccination (<2 months), or symptoms of a respiratory tract infection or common cold within the past two weeks were asked to participate at a later stage. Additional questionnaires were taken to further assess frailty, including a KATZ-ADL^34^ questionnaire for daily activities, a KATZ-iADL^35^ questionnaire for the daily use of instruments and a 6-cognitve Impairment Test (6-CIT). Additionally, handgrip strength was measured. Ethical approval was obtained from the Medical Ethical Committee Leiden-Den Haag-Delft (NL77841.058.21).

The TINO study design included two scheduled visits, with half of the participants in each group undergoing an additional sampling visit three months later. In cases where an acute respiratory infection was suspected during those 3 months, participants were instructed to contact the study team. Additional sampling was performed when a respiratory pathogen was detected by PCR through routine diagnostics at the LUMC. These follow-up samples were collected within two weeks of symptom onset, and again at 1, 3, and 5 months after symptom onset.

Samples obtained from the URT by nasal curettage from SARS-CoV-2 infected patients (acute phase, mild and moderate disease) were additionally used from the BEAT-COVID and SARS-RESPONSE studies. Ethical approval for these studies was obtained from the Medical Ethical Committee Leiden-Den Haag-Delft (NL73740.058.20). The trial was registered in the Dutch Trial Registry (NL-OMON24946).

#### Sample collection and cell isolation

Nasal-derived cells collected from the inferior turbinate using curettage were isolated as described previously^36^. Briefly, curettes (ASL Rhino-Pro©, Arlington Scientific) were used to scrape cells from the inferior turbinate. Two curettes per nostril were used and stored in a 15 mL falcon tube placed on ice containing phosphate-buffered saline (PBS), 0.5% heat-inactivated fetal calf serum (hiFCS, PAN Biotech, United Kingdom), and 5 mM ethylenediaminetetraacetic acid (EDTA). Cells were dislodged from the curettes by repeated pipetting with collection medium from the same tube. Cells were spun down (450g for 5 minutes at room temperature) and pellets from the two 15 mL Falcon tubes (left and right nostril) were pooled, resuspended in 1 mL Cryostor CS10 (Stemcell) and aliquoted into two cryovials. Step-wise freezing was performed by using pre-chilled Mr. Frosties prior long-term cryopreservation in a liquid nitrogen tank.

Nasal lining fluid was collected from the inferior turbinate using 7.0mm synthetic absorptive matrixes (Nasosorption^TM^ FX-I, Hunt developments). The filter strip of the nasosorptions were inserted fully in the nose, and the same nostril was closed with the index finger for 60 seconds. Collected samples were transported back to the laboratories on ice and stored at - 70°C until elution. Samples were eluted in gender matched batches with equal proportions of young, healthy older adults and frail older adult samples, including one mock elution of an untouched nasosorption. Nasosorption samples were eluted by adding 100 µL of sterilized elution buffer (PBS + 1% high-grade sterile BSA + 0.05% Triton-X) dropwise on both sides of the absorptive matrix. The overflow was then aspirated and re-added to the nasosorption device, until the matrix was visibly saturated. The nasosorptions were then centrifuged at 4,000g for 10 minutes at 4°C. Total volume of eluted sample was recorded and the eluted sample was resuspended and moved to a sterile Eppendorf tube, and spun at 16,000 g for 10 minutes at 4°C, to pellet bacteria. The supernatant volume was kept at −70°C until further procedures. All work was performed in sterile conditions to avoid sample contamination.

Peripheral blood mononuclear cells (PBMCs) were obtained by standard Ficoll-Isopaque separation of sodium-heparin peripheral blood collection tubes and 5-10×10^6^ cells were aliquoted per cryotube in freezing medium (RPMI 1640 Medium (Gibco, United Kingdom) containing 100 µg/mL streptomycin, 100 U/mL penicillin, 10% hiFCS and 10% DMSO and stored in the vapor phase of liquid nitrogen. Plasma was obtained from 2 mL EDTA peripheral blood collection tubes by centrifuging the tubes at 400g for 7 minutes. Plasma was then taken and centrifuged again at 600g for 7 minutes. Plasma was cryopreserved in cryotubes at −70°C until further procedures.

#### Nasal phenotyping using spectral flow cytometry

Cryopreserved nasal-derived cells were thawed in three separate batches in a 37°C water bath and transferred to a 15 mL falcon tube and washed by slowly adding 4 mL pre-warmed thawing medium (RPMI 1640 Medium, containing 20% hiFCS, 100 µg/mL streptomycin, and 100 U/mL penicillin). Sample selection was performed to achieve the closest possible matching for age and sex with equal distribution across batches . Samples were centrifuged for 7 minutes at 400g and supernatant was discarded. Thawed nasal samples were then taken up in 180 µl PBS and transferred to a V-bottom 96-well Nunclon™ Delta Surface plate (Thermo Fisher Scientific, Denmark), and cells were spun down for 7 minutes at 400g. After centrifuging and removing supernatant, cells were resuspended and stained in 50 µl for 15 minutes at RT with LIVE/DEAD™ Blue fixable Blue Dead Cell Stain (Invitrogen, USA) solution at 1:500 dilution mixed with anti-human Fc Receptor (FcR) Binding Inhibitor (Invitrogen, USA). Then extracellular staining was performed by adding 50 µl of 2x concentrated mix containing a large phenotyping panel (n=33) of monoclonal antibodies directed against extracellular proteins (**Supplementary table 2**) prepared in FACS buffer (PBS containing 2 % (v/v) BSA, 2 mM EDTA) with 10 % (v/v) BD Horizon™ Brilliant Stain Buffer Plus (BD Biosciences, USA) for 15 minutes at room temperature (RT). After extracellular staining, cells were washed twice with FACS buffer and then resuspended in 200 µL of FACS buffer for acquisition. Cells were acquired with fluidics boost on a 5-laser Aurora Cytometer (Cytek Biosciences, Inc, USA). Reference controls were made using either AbC™ Total Antibody Compensation Beads (Invitrogen, USA) or PBMCs. An additional unstained control, gated on monocytes (FSC/SSC) was used for unmixing. After acquisition, epithelial cells, granulocytes (CD45+CD66b+) and live lymphocytes (CD45+CD66b-) were gated using Flowjo v10 and exported (**Supplementary Fig. 3A**). Exported live lymphocyte fcs files were uploaded in OMIQ software (omiq.ai) for batch correction by sub clustering each batch into 5 major clusters using FLOWsom k=5 with a dimension of 15×15 and 25 training iterations, followed by Cytonorm with 101 quantiles to apply normalization across batches. Samples with less than 50 lymphocytes were discarded. Supervised clustering was then performed by manual gating in OMIQ (**Supplementary Fig. 3B).** Unsupervised clustering was performed on normalized live lymphocytes (CD45+CD66b-) using Elbow meta clustering using FLOWsom with a dimension of 15×15 and 50 training iterations. Immune cell numbers were normalized to epithelial cell numbers within each sample, enabling an independent assessment of immune cell populations while accounting for variation in sample yield^16^. Samples in which no cells were detected for a given population were excluded from analyses involving that specific population.

#### Cytometry by Time Of Flight (CyTOF) staining

Nasal-derived samples from individuals with acute SARS-CoV-2 infection in the TINO study were barcoded and measured in one single batch of twenty samples as described previously^20^. One aliquot from a reference sample was included to normalize staining against previously acquired batches^16, 20^. Nasal cells were thawed similarly as described above, but 50% hiFCS was used instead. Staining was performed as described previously. In short, for each batch, nasal-derived cells and reference PBMCs were resuspended in 50 µl of Perm/Wash, and 50 µl of barcode mix targeting β2-microglobulin (B2M) was added to each individual sample in a 6-choose-3 scheme using cadmiums 106, 110, 111, 112, 114 and 116. Samples were incubated for 30 minutes at RT and then washed with 4 mL of Cell Staining Buffer (Fluidigm). Cells were centrifuged for 5 minutes at 800g, and the supernatant was removed, resuspended and combined into 3 mL of Perm/Wash. Cells were centrifuged again for 5 minutes at 800g and were resuspended in 45 µl of Perm/Wash. FcR block (Biolegend; 5 µl) and sodium heparin (0.5 µl, 100U per mL) were added to prevent aspecific binding of antibodies, and cells were incubated for 20 minutes at RT. Then, 50µl of antibody cocktail (**Supplementary Table 3**) was added, followed by a 45 minute incubation at RT. Cells were then washed twice with 2 mL of Cell Staining Buffer and centrifuged for 5 minutes at 800g. DNA was then stained overnight at 4°C using 1 mL of Fix and Perm buffer (Fluidigm) containing 1,000 fold diluted Intercalator-Ir (Fluidigm). Cells were then washed with Cell Staining Buffer, counted and divided into tubes of 1×10^6^ cells and pelleted. Pellets were washed and resuspended in cell acquisition solution (CAS, Fluidigm) with EQ Four Element Calibration Beads (Fluidigm) and acquired on a Helios mass cytometer (Fluidigm) with CyTOF Software (v7.0.8493) at the Flow Cytometry Core Facility (FCF) of LUMC in Leiden, the Netherlands.

### CyTOF analysis

Compensation, filtering for debris and normalization beads was performed as described before using “CyTOFclean” and “CATALYST” packages^20^. Correct compensation of data was visually checked with one by one plots. Epithelial cells and immune cells were manually gated and exported as .fcs files based on DNA dye, CD45 and EpCAM expression, with exclusion of cPARP positive apoptotic cells, as well as immune doublets (CD14+CD3+ , CD66b+CD3+ , CD14+CD66b+). Nasal-derived cells with less than 100 immune and/or less than 100 epithelial cells were removed, as they indicated unsuccessful sampling.

Samples from this study were integrated with previously published cohorts acquired using the same panels and reference control, with the exception of CD9 and CD20 which were omitted from the first batches measured. We combined 46 previously acquired samples from SARS-CoV-2 infected individuals with 18 additional infected individuals (10 of which from the TINO study and 8 from the BEAT-COVID study). In addition, 25 previously acquired samples from uninfected individuals were included. All 65 infected individuals were Europeans and within 26 days of infection onset (median 11 days, IQR 8-16 days). Of these infected individuals, 32 had a mild infection and 33 patients were hospital admitted due to hypoxia (moderate infection) and no intensive care unit admitted patients were included (**Supplementary table 4**). Cellular immune cell data (.fcs) from these samples were uploaded in OMIQ software for batch correction. Granulocytes (CD45^+^CD66b^+^) were gated out and remaining lymphocytes were further used for batch normalization. For batch normalization, lymphocytes were sub clustered using all markers except CD19/CD20, CD9 and EpCAM into 5 clusters using FLOWsom k=5 with a dimension of 15×15 and 25 training iterations, followed by Cytonorm with 101 quantiles to apply normalization across batches. We manually gated CD4^+^ and CD8^+^ Tem (CD45RA^-^CCR7^-^) and Trm (CD69^+^) cells (**Supplementary table 4** and **Supplementary Fig. 6**). Samples in which no cells were detected for a given population were excluded from analyses involving that specific population.

### *In vitro* co-culture of autologous PBMCs in a 3D nasal epithelial model

A detailed step-by-step protocol for the *in vitro* 3D nasal epithelial co-culture model with PBMCs has been published previously^21^. In short, nasal epithelial organoids were generated from primary nasal epithelial cells collected from TINO participants using nasal curettage as described above. Primary nasal epithelial cells were treated twice with TripLE Express (Thermo Fisher Scientific, USA) for 7 minutes at 37°C. Stop medium (RPMI 1640 medium containing 100 µg/mL streptomycin, 100 U/mL penicillin and 10% hiFCS) was added and single cell suspension was centrifuged at 230g for 7 minutes. Cells were then seeded in 30 µl Cultrex Reduced Growth Factor Basement Membrane Extract, Type 2, Select (BME-2, Bio-Techne, Ireland) droplets. Primary epithelial cells were cultured at 37°C and 5% CO2 in airway epithelial medium composed of Advanced Dulbecco’s Modified Eagle Medium (DMEM)/F-12 medium (Thermo Fisher Scientific, USA) supplemented with 10 mM HEPES (Thermo Fisher Scientific, USA), 1x GlutaMAX (Thermo Fisher Scientific, USA), 100 μg/mL primocin (InvivoGen, USA), 25 ng/mL R-spondin-1 (Thermo Fisher Scientific, USA), 25 ng/mL FGF-7 (Thermo Fisher Scientific, USA), 100 ng/mL FGF-10 (Miltenyi, Germany), 100 ng/mL noggin (Thermo Fisher Scientific, USA), 500 nM A83-01 (Selleckchem, USA), 5 μM Y-27632 (Cayman Chemical, USA), 500 nM SB202190 (Merck, Germany), 1x B27 supplement (Thermo Fisher Scientific, USA), 1.25 mM n-acetylcysteine (Merck, Germany) and 5 mM nicotinamide (Merck, Germany). Airway epithelial medium was refreshed every 3-4 days. After two weeks of culture, organoids from young adults, healthy fit or frail older adults were harvested from BME-2 droplets using cold PBS and co-cultured in the presence or absence of freshly thawed autologous PBMCs in 96-well flat-bottom plates (per condition: 20.000 epithelial cells and 100.000 PBMCs) in T cell medium (Iscove’s Modified Dulbecco’s Medium (IMDM, Capricorn Scientific, Germany) supplemented with 100 IU/mL penicillin (Eureco-Pharma, Netherlands), 100 μg/mL streptomycin (Sigma-Aldrich, Canada), 2 mM L-glutamin (Sigma-Aldrich, Canada), 50 IU/mL IL-2 (R&D Systems, USA), 5% hiFCS and 5% heat-inactivated human serum (Sanquin, Netherlands)) at 37°C and 5% CO2. For TGF-β blocking experiments, a TGF-β inhibitor (10 μg/mL, SB431542, Selleck Chemicals, USA) was weekly added to the co-culture model. Co-cultures were maintained for up to 28 days, during which 25% of the medium was refreshed every 2-3 days with new T cell medium. Supernatants collected on days 7, 14, 21 and 28 were stored at −20°C for subsequent cytokine analysis.

To assess the induction of Trm cells in the co-culture, cells were passed through a 40 μm strainer and transferred to a new 96-well V-bottom plate to separate epithelial cells in the 3D NEM from immune cells. Cells were washed twice with PBS and stained in 100 μL PBS supplemented with LIVE/DEAD™ Blue fixable Blue Dead Cell Stain solution at 1:500 dilution for 15 minutes at RT. After washing the cells with FACS buffer and centrifugation (230g, 7 minutes), cells were stained in 100 μL extracellular antibody mix (**Supplementary Table 5**) for 15 minutes at RT. Cells were again washed with FACS buffer, resuspended in 150 μL FACS buffer and acquired on a 3-laser Aurora Cytometer (Cytek Biosciences, Inc, USA). Reference controls were prepared using either Anti-Mouse Ig, κ/Negative Control Compensation Beads (BD, USA) or PBMCs. An unstained control was used for every timepoint.

#### *In vitro* induction of tissue-resident memory T cells by TGF-**β**1 and IL-15

PBMCs from either young adults, healthy older adults or frail older adults were thawed and cultured at 37°C and 5% CO2 in 96-well flat-bottom plates (per condition: 100.000 cells) in T cell medium. PBMCs were stimulated at initiation of culture and after 7 days with 0.5, 5.0 or 20.0 ng/ml TGF-β1 (Thermo Fisher Scientific, USA) and/ or 10.0 ng/ml IL-15 (Thermo Fisher Scientific, USA). Every 2-3 days 25% of medium was refreshed with new T cell medium. PBMCs were harvested at day 14 to assess the Trm phenotype within the CD8^+^ T cell population using spectral flow cytometry (**Supplementary Table 5**), similarly as described above.

#### ELISA

Culture supernatants were collected from 3D NEMs following 14 days of culture. TGF-β1 and IL-15 production by nasal epithelial cells were quantified in the collected culture supernatants using standard enzyme-linked immunosorbent assays (ELISA) according to the manufacturer’s instructions (TGF-β1 kit: Bio-Techne, USA and IL-15 kit: BD, USA).

#### Olink proteomics

Detection of inflammation-related proteins from nasal lining fluid and EDTA plasma, was performed using the Olink Target 96 Inflammation panel (Thermo Fisher) and was measured on a Biomark^TM^ HD90602 PCR machine (Standard BioTools) at Leiden, Leiden University Center for Infectious Diseases, Leiden, The Netherlands. Before samples were ran, the integrated fluidic circuit was primed on a IFC Controller HX (Standard BioTools). The NPX Signature software version 1.17.0 was used for QC analyses and to generate normalized protein expression (NPX) values at a log2 scale. In total, 86 out of 88 nasal lining samples and 83 out of 88 EDTA plasma samples passed the quality control and were analyzed. Nasal lining data was normalized by background subtraction of the control sample consisting of only elution buffer to define background values per protein. Sixteen proteins were below the negative control and were taken out for further analysis.

#### Single cell RNA sequencing

Cryopreserved nasal-derived cells were thawed in two batches in a 37°C water bath and transferred to a 15 mL falcon tube and washed by slowly adding 4 mL pre-warmed thawing medium, similarly as above, containing 50% hiFCS. Samples were centrifuged for 7 minutes at 400g at 4°C and supernatant was discarded. For the first batch, 12 samples were pooled on ice by resuspending the pellets in 1 mL RPMI 1640 Medium, containing 10% hiFCS, 100 µg/mL streptomycin, and 100 U/mL penicillin. The pooled cell-suspension was then filtered using a 40µm cell strainer and centrifuged for 7 minutes at 400g at 4°C. Dead cells were removed with the dead-cell removal kit MojoSort™ (Biolegend). In summary, the pellet was taken up in 94 µL of Mojosort Buffer, 1 µL of RNAse OUT^TM^ (Invitrogen, USA cat.10777019) and 5 µL of MojoSort human dead-cell removal beads and incubated for 15 minutes in the fridge. The cell-suspension was then taken up in 3 mL Mojosort Buffer and transferred to a 5 mL Falcon round bottom polypropylene tube (Corning, Mexico) and placed inside the 5ml MojoSort tube magnet (Biolegend) and incubated for 3 minutes on ice. The supernatant was then collected in a 15 mL Falcon tube and another 2 mL was added to the 5 mL Falcon round bottom tube, incubated for 3 minutes and subsequently collected and added to the 15 mL Falcon tube. The enriched fraction was then centrifuged at 450g for 7 minutes at 4°C. Cell pellet was taken up in 25 µL of PBS containing 2% BSA and 1 µL RNAse OUT^TM^ (Invitrogen) and transferred to a loBind, RNAse free 1.5 mL Eppendorf tube. The second batch containing 18 samples was processed similarly but each sample was incubated with 1/100 TotalSeq™-C Hashtags (Biolegend) after thawing for 20 minutes on ice (**Supplementary Table 6**). Samples were washed with 5 mL FACS buffer and centrifuged at 400g for 7 minutes at 4°C. Samples were then resuspended in 500 µL of FACS buffer and pooled in a 15 mL tube containing 3 mL FACS buffer already. The pooled multiplexed cell-suspension was then filtered and enriched for live cells similarly as above. Library preparation was carried out using the Chromium GEM-X Single Cell 5’ RNA Library Kit on a GEM-X chip (∼60,000 cells per lane) with barcoded beads according manufacturer’s protocol. Libraries were prepared for (i) 5′ gene expression (transcriptome), (ii) T cell receptor (TCR) V(D)J enrichment, and (iii) hashtag oligonucleotides (HTO; hashtag library) using the 10x Feature Barcoding/HTO workflow. The HTOs were only prepared for batch 2 and 3. The following changes were made for the preparation of the T cell receptor library: an input of 4 µL of cDNA was used and 2 additional PCR cycles were implemented. All libraries were quantified (Qubit) and evaluated for size distribution (Bioanalyzer) prior to pooling. Libraries were sequenced on an Illumina NovaSeq X 10B.

#### Bulk whole-transcriptome sequencing

Bulk whole transcriptome sequencing (WES) was performed on 1×10^6^ lysed PBMCs in order to demultiplex batch 1 of single cell RNA (scRNA) sequenced nasal-derived cells. These individuals were mapped by using a single-nucleotide polymorphism (SNP) approach, see below. The NEBNext Ultra II Directional RNA Library Prep Kit for Illumina was used to isolate RNA from the PBMC samples, according the protocol “NEBNext Ultra II Directional RNA Library Prep Kit for Illumina” (NEB cat#E7760S/L). Briefly, rRNA was depleted from total RNA using the Qiagen fast select kit (cat#334387). After fragmentation of the rRNA reduced RNA, a cDNA synthesis was performed. This was used for ligation with the sequencing adapters and PCR amplification of the resulting product. The concentrations of the samples after library preparation were determined using a fluorescence-based assay, and the quality was assessed using the QIAxcel Connect system. All samples passed QC and sequencing using the NovaSeqX plus was performed according to manufacturer’s protocols. A final loading concentration of ∼ 160 pM of DNA was used.

### Single cell RNA-Seq analysis pipeline

#### SNP-based de-multiplexing

Cells were clustered by subject identity using a modified version of *vireo*, which infers sample membership from genetic variation in scRNA-sequencing data. In its original formulation, *vireo* maximizes the joint probability of the observed allelic counts together with: ,,The likelihood of those counts given the latent cell-to-subject assignments and the latent subject genotypes”, and ,,The prior probabilities on both the subject genotypes and the cell-to-subject assignments”. The iterative process of maximizing the joint probability allows to infer the latent variables of interest. The original ‘vireò method has limited options for specification of the prior probabilities for cell to subject assignment. We have reimplemented the method and modified the prior part to allow for more flexible specification. We applied the modified method to assign cells to subjects in our dataset. As the prior probabilities of the genotypes of the subjects we used the genotype predictions calculated from the WES data of the individual subjects. First, based on the WES data we selected the variant positions that were identified as heterozygous or homozygous alternative in at least one subject. Then, we calculated the total number of reads observed at these positions in the mixed-subject (pooled) scRNA-seq data. We selected 3000 positions with the highest read coverage in the scRNA data. In our modified method we can specify a non-uniform cell to subject assignment prior.

This prior utilizes the known sex of the subjects (6 females, 6 males for batch 1). For each cell, we calculated the likelihoods that it originated from a female or a male, based on the observed number of cell reads mapping to the Y chromosome relative to the total number of mapped reads of that cell. These probabilities were then used as the cell-to-subject assignment prior in the modified method. With the above specifications, we applied the modified method to assign cells to 12 subjects. We ran the method with 200 randomly chosen initializations with a maximum of 200 iterations per initialization, and selected the iteration with the highest ELBO (Evidence Lower Bound). When ELBO did not improve significantly for 5 consecutive iterations, we stopped the optimization process. Each cell was labeled according to their sample ID, and only mapped cells were extracted and integrated with other batches.

#### Hashtag-based de-multiplexing

Single-cell RNA sequencing data generated using the 10x Genomics platform were processed in R (v4.4.1), using the Seurat package (v5.4.0). Gene expression and HTO count matrices were imported from Cell Ranger output using the Read10X function. Filtered feature-barcode matrices were used for downstream analysis. Gene expression counts were used to create a Seurat object, retaining genes expressed in at least three cells, HTOs were extracted and stored as a separate assay within the Seurat object, enabling downstream demultiplexing analysis. The resulting multi-assay Seurat object, containing both RNA and HTO modalities, was saved for subsequent preprocessing and quality control steps. Ambient HTO signal was estimated using raw 10x data. Empty droplets were identified using the emptyDrops method, and barcodes not classified as cells were used to define the background HTO profile. Background droplets were used to build a Seurat object representing ambient signal. HTO counts from these droplets were summed and normalized to obtain the relative abundance of each HTO tag in the ambient background. HTO-based demultiplexing was then performed using HTODemux (Seurat v5.4.0) and hashedDrops (DropletUtils, v1.24.0). Each method classified cells as singlets, doublets, or negatives based on HTO count distributions. For hashedDrops, an ambient HTO profile derived from background droplets was incorporated to improve classification accuracy. Final cell identities were determined by combining results from both methods using a consensus approach (union or intersection), guided by classification agreement (**Supplementary Fig.17**). RNA quality control (QC) was performed using standard single-cell metrics, including the number of detected genes, total UMI counts, and the percentage of mitochondrial gene expression. Additional metrics, such as gene complexity (log10 genes per UMI), were used to identify low-quality cells. Cells were filtered using median absolute deviation (MAD)-based thresholds applied to library size, gene counts, mitochondrial percentage, and complexity. Cells that failed QC criteria, identified as doublets or unassigned by demultiplexing, were excluded from downstream analyses. The final filtered datasets were retained for downstream processing and comparison of processing strategies. Cells classified as doublets by both methods were labeled as high-confidence doublets. For non-doublet cells, hashedDrops assignments were prioritized, with HTODemux used to fill missing labels. Cells classified as negative or unassigned were excluded (**Supplementary Fig.17**). The filtered dataset was processed in Seurat using SCTransform normalization with regression of mitochondrial content, followed by PCA, UMAP visualization, and graph-based clustering. A k-nearest neighbor purity filter (RANN package, v2.6.2) was applied in HTO space, and cells with neighborhood purity < 0.5 were removed. Cell types were annotated using canonical marker genes, and epithelial (EPCAM[) and immune (PTPRC[) compartments were defined for downstream analyses.

### Integration and analysis

Single-cell datasets from the two HTO-demultiplexed batches were merged with the SNP-demultiplexed batch, and batch effects were corrected using Harmony (Version 1.2.4). Clusters were identified using the Harmony-corrected embeddings and visualized across batches to assess integration quality (**Supplementary Fig. 17g**). Cell types were further reviewed and annotated based on canonical marker gene expression. To map different T cell populations in greater detail, clusters enriched for T lymphocytes were subsetted from the globally integrated dataset. The subsetted data were reprocessed using the SCTransform normalization workflow, and highly variable genes were reidentified. T cell receptor (TCR) genes (TRAV, TRBV, TRDV, and TRGV families) were excluded from the variable feature set to prevent clustering driven by clonal variation. Principal component analysis was performed, and batch effects within the T cell subset were corrected using Harmony. The batch-corrected embeddings were used to construct a shared nearest neighbor graph, followed by graph-based clustering across multiple resolutions and visualization using Uniform Manifold Approximation and Projection (UMAP). Cluster marker genes were identified using the FindAllMarkers function in Seurat with a minimum expression threshold of 25% of cells and a log fold-change threshold of 0.25. Mitochondrial, ribosomal, and Ensembl identifier genes were excluded from downstream marker analyses. Cluster distributions across sequencing batches were assessed by calculating cell counts and proportions per cluster.

#### T cell receptor sequencing

TCRαβ sequences of T cell populations were identified as previously described^37, 38^. In short, total RNA (10 μL) was extracted using the ReliaPrep RNA cell Miniprep system (Promega) from cryopreserved (Cryostor CS10) nasal samples collected using nasal curretage longitudinally 3 months apart or 1, 3 and 5 months after SARS-CoV-2 infection. Additionally, PBMCs were stimulated with overlapping peptides for SARS-CoV-2 and activated T cells were separately sorted for CD4 and CD8 T cells and RNA was similarly extracted.

Preamplification of the cDNA was performed on samples containing RNA from 500 or fewer cells. Barcoded TCR PCR product was generated in two rounds of PCR. In the second PCR, the first purified PCR product was used to include a two-sided six-nucleotide barcode sequence that allows for discrimination between TCRs of different T cell populations. PCR products of different T cell populations were pooled, after which TCR sequences were identified by NovaSeq (GenomeScan). NovaSeq data were analyzed using MiXCR software (v3.0.13) to determine the Vα and Vβ family and CDR3 regions using annotation to the IMGT library (http://www.imgt.org; v6). CDR3 sequences with a frequency lower than 0.01% of total reads, those that contained a stop codon, did not start with cysteine, or occurred in all samples were excluded from analysis. The frequencies of overlapping CDR3 amino-acid (AA) sequences between two timepoints were calculated using depth-corrected overlap using the following formula whereby the number of overlapping clones between two timepoints were normalized to the lowest number of clones identified at either timepoint. Ct1 and Ct2 represent the number of unique CDR3 AA sequences and Ct1[Ct2 represents the number of shared sequences.

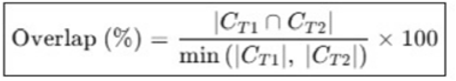

## Data availability

Raw datasets, including TCR sequencing, CyTOF, spectral flow cytometry, and single-cell RNA sequencing data, have been deposited in the Zenodo repository and are available at https://doi.org/10.5281/zenodo.xxxx. Researchers may request access by submitting a research proposal or protocol to the corresponding contact person. Following approval, the data will be shared without restrictions. Access is controlled due to the possibility that clinical data, despite anonymization, may still be re-identifiable. No fees will be charged for data access.

## Code availability

All packages, functions and key parameters used for analyses have been included the Methods section. Scripts used are deposited in GitHub at https://github.com/spjochems/TINO_Tcell.

## Contributions

W.H. contributed to Conceptualization, Project administration, Investigation, Methodology, Software, Formal analysis, Data curation, Visualization, Writing - original draft, and Writing - review & editing. L.A.K. contributed to Investigation, Formal analysis, and Visualization. Y.S. contributed to Data curation, Software, Formal analysis, and Methodology. A.C.d.K., R.A.M.S., and I.v.d.V. contributed to Investigation and Resources. R.S.H. contributed to Investigation. E.J.d.M. contributed to Investigation and Data curation. M.K. contributed to Investigation. T.R., G.E.L.S.S., M.T.W., and M.v.d.E. contributed to Resources and Investigation. S.L.K. contributed to Supervision and Data curation. J.J.C.d.V. contributed to Investigation and Resources. A.H.E.R. contributed to Resources and Investigation. S.M.K. contributed to Formal analysis and Data curation. C.E.W.S. and D.O.M.K. contributed to Resources. M.Y. contributed to Supervision. A.M.v.d.D. and P.S.H. contributed to Supervision. M.H.M.H. contributed to Supervision. S.P.M. and G.H.G. contributed to Resources, Supervision, and Project administration. S.P.J. contributed to Conceptualization, Supervision, Formal analysis, Software, Visualization, Writing - original draft, and Writing - review & editing, Project administration, and Funding acquisition.

## Acknowledgements

We thank all research nurses and physicians responsible for sample collection, and we are especially grateful to all healthy volunteers for their participation in this study. We further acknowledge general practise “Poelpolder”, and general practise “De Lindehoeve” as part of the Extramural LUMC Academic Network (ELAN) for their essential contributions to participant recruitment. The authors gratefully acknowledge the flow cytometry core facility (FCF) at LUMC, Leiden, the Netherlands for technical support regarding spectral flow cytometry and CyTOF. This work was supported by an NWO grant (OCENW.KLEIN.461), a ZonMw grant (10430072110011) and a LUMC Gisela Thier Fellowship award (2020) to SPJ.

## Supplementary Figures

**Supplementary Figure 1.**
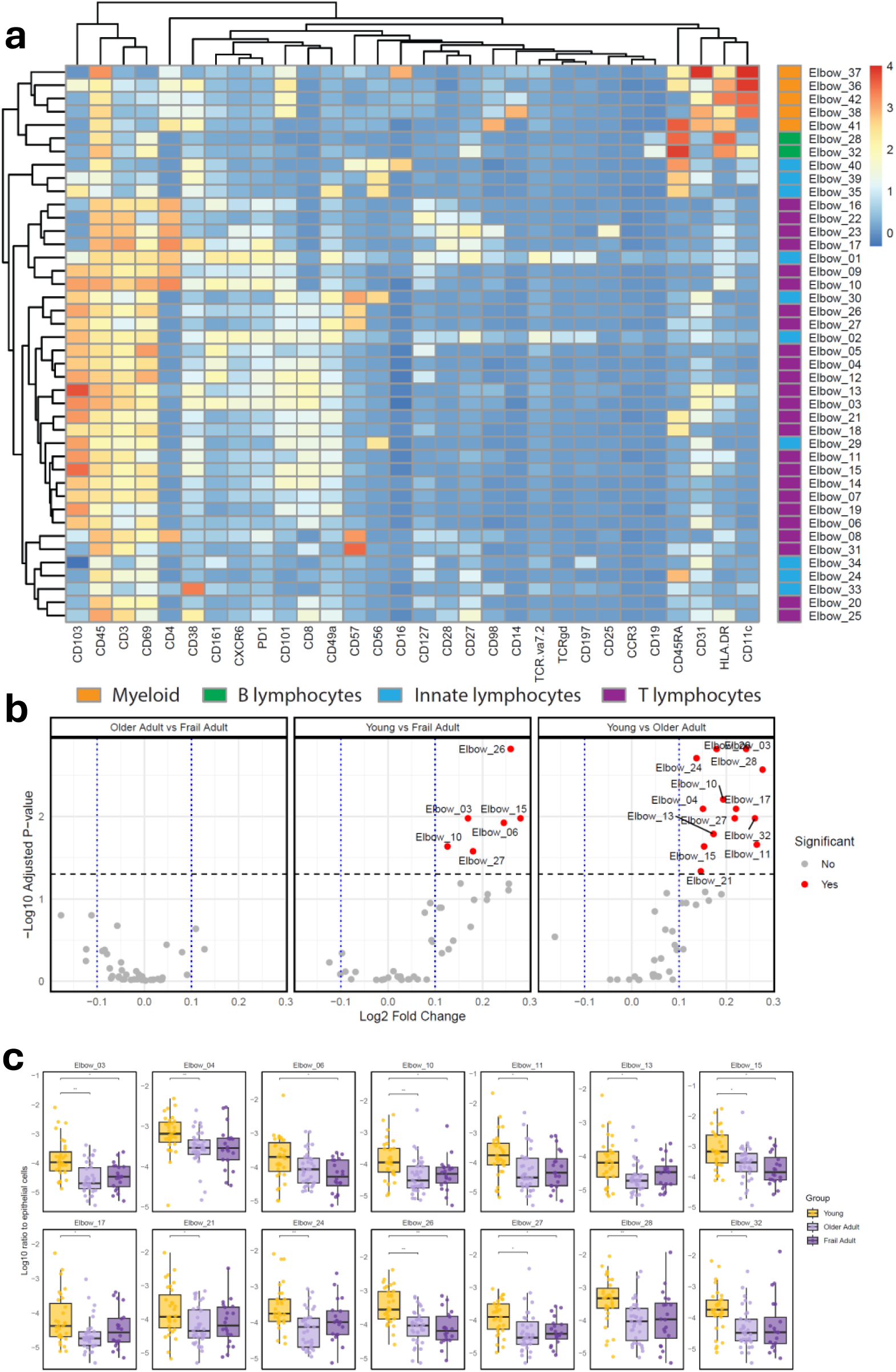
Defining immune cell subsets using unsupervised clustering and performing group comparisons. **a)** Heatmap showing the median expression of different markers (x-axis) for unsupervised clustered populations using FLOWsom Elbow algorithm (n=42, y-axis). Rows and columns were hierarchically clustered using Euclidean distance and complete linkage. **b**)Volcano plots depicting differential abundance of FLOWsom Elbow clusters between young and older adults, young and frail adults, and older and frail adults. Each dot represents one immune cell population. The dashed horizontal line indicates the significance threshold (adjusted P < 0.05). Red dots represent significantly different populations (FDR < 0.05), whereas grey dots indicate non-significant populations. **c**) Boxplots of Elbow clusters that were differentially abundant between groups, expressed as the log10 ratio of immune cells to epithelial cells. Statistical differences were assessed with Kruskal Wallis tests followed by Wilcoxon signed rank tests with Benjamini-Hochberg correction for multiple testing. \**P* < 0.05; ** *P < 0.01*, *** *P < 0.001*

**Supplementary Figure 2.**
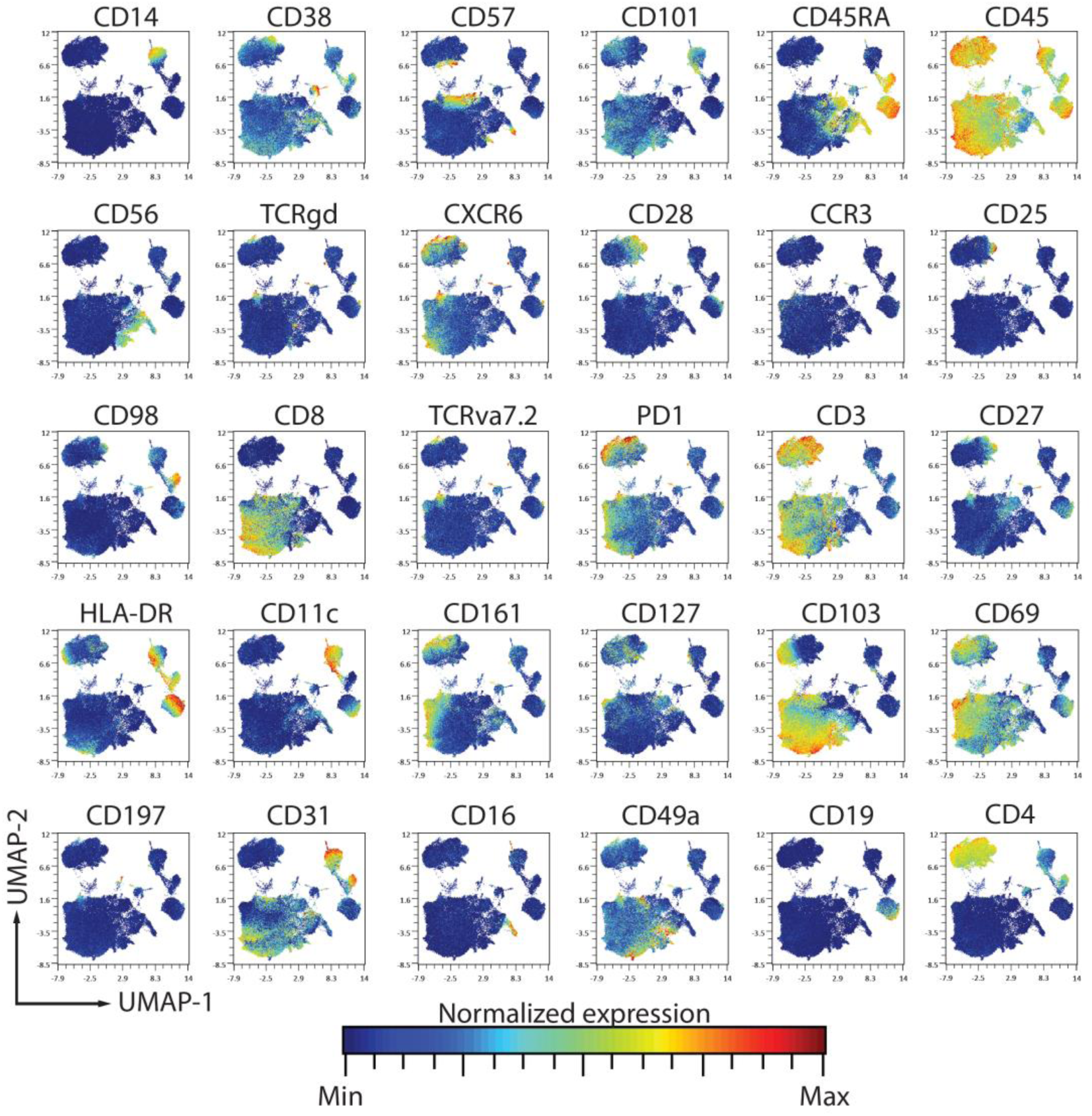
Uniform Manifold Approximation and Projection (UMAP) of all nasal-derived immune cells and marker expression. UMAPs of 102910 nasal-derived immune cells (excluding granculocytes) are shown and overlaid with the normalized expression of markers that were used to perform unsupervised clustering.

**Supplementary Figure 3.**
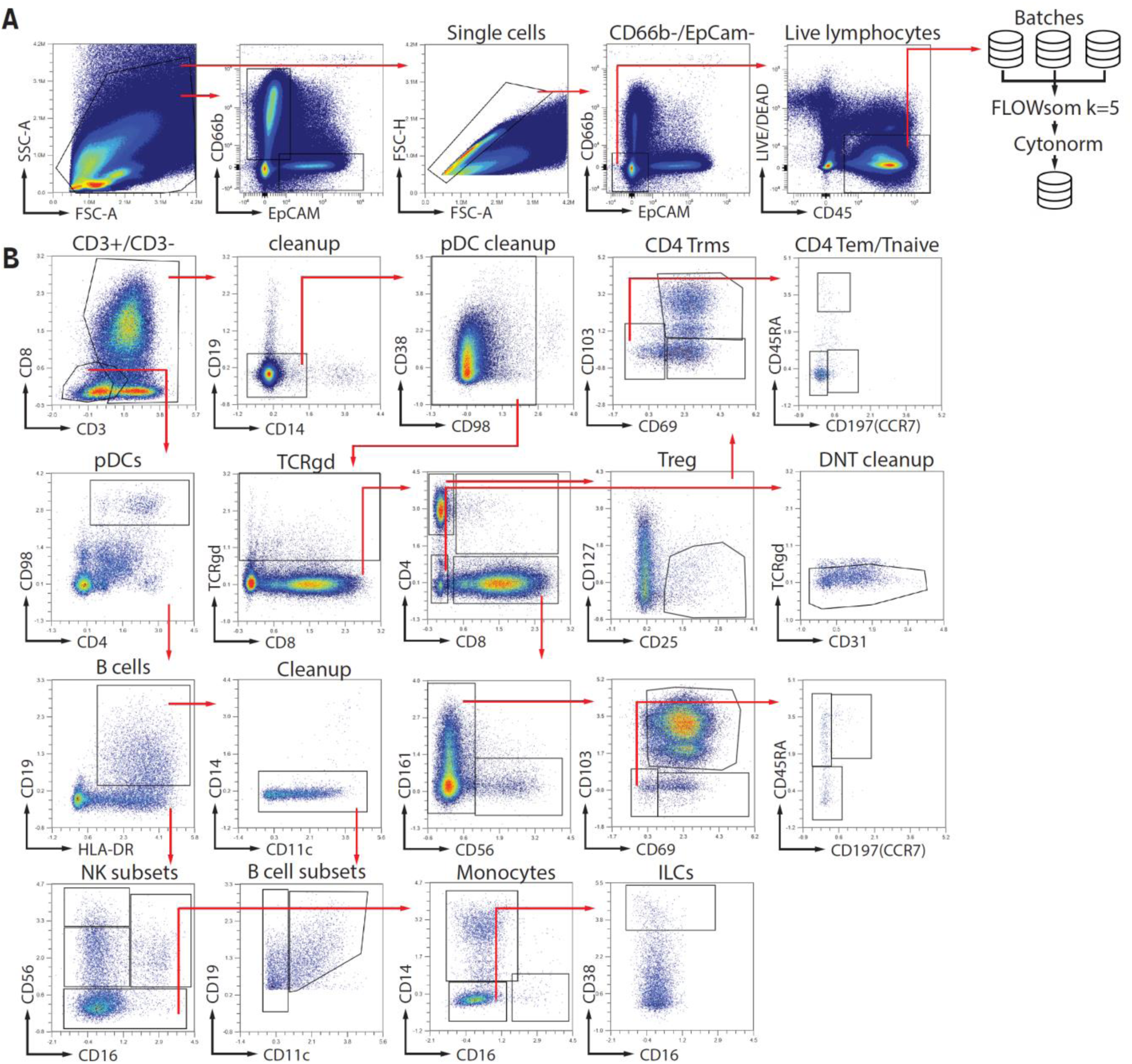
Gating strategy of phenotyping nasal-derived immune cells. Cryopreserved nasal samples were thawed, stained and acquired in 3 batches on a 5-laser Aurora. **A**) Live lymphocytes, excluding granulocytes (CD45+CD66b-), granulocytes and epithelial cells were first gated and exported. Batch correction using FLOWsom k=5 and Cytonorm was then applied for all markers except CD66b, Epcam, CD45 and Live-dead for live lymphocytes. **B**) Normalized live lymphocytes were used to further gate different immune cell populations.

**Supplementary Figure 4.**
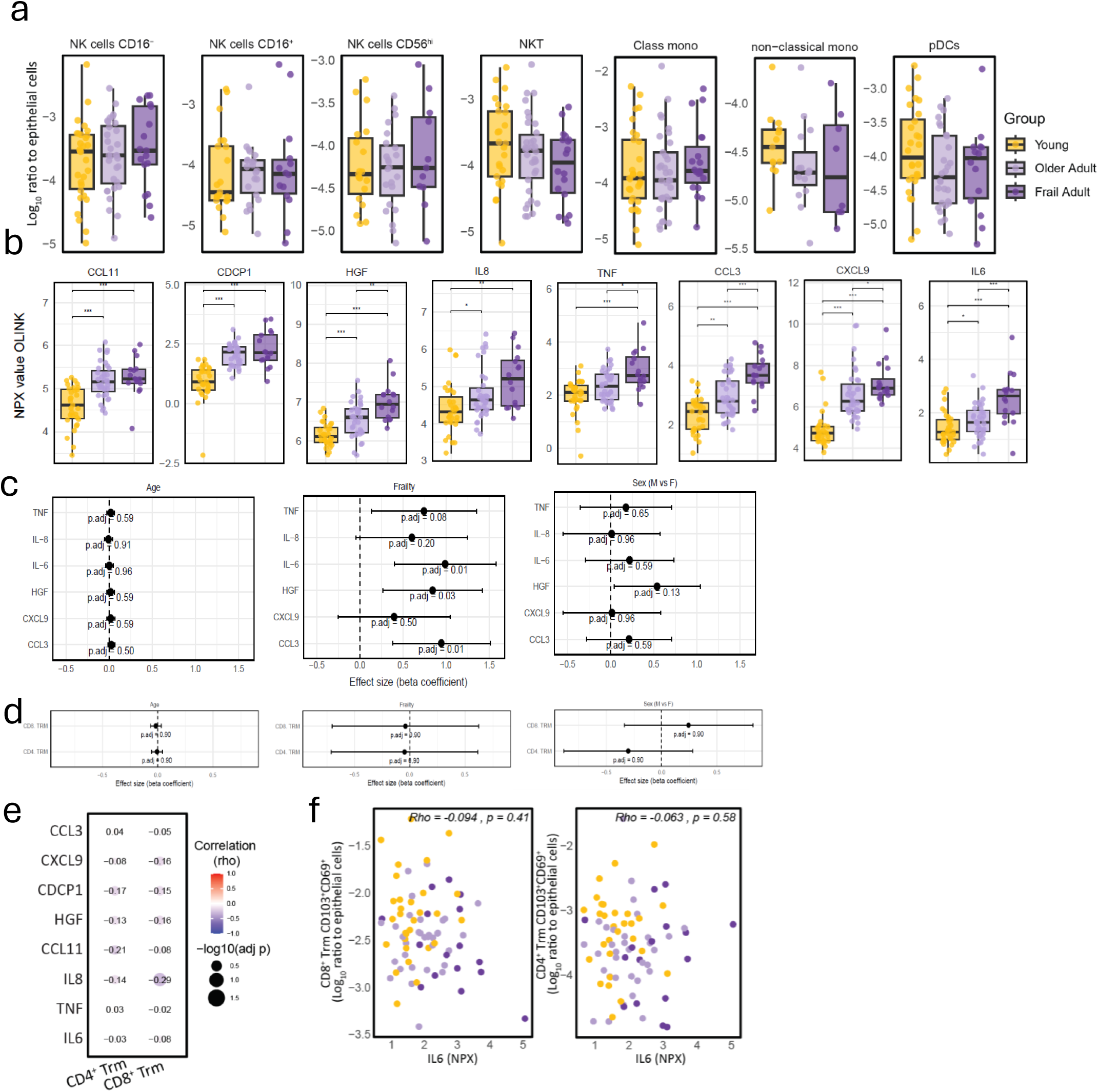
Stable immune cell populations in the upper-respiratory tract across age. Nasal cells were collected by curettage (RhinoPro©) and cryopreserved prior to analysis. Immune cell populations were manually gated and normalized to the number of epithelial cells, expressed as the log10 ratio of immune cells to epithelial cells from the same sample. **a**) manually gated innate immune cell populations expressed as the log10 ratio of immune cells to epithelial cells. Data are presented as boxplots and compared between healthy young adults, healthy older adults, and frail older adults. **b**) Shown are inflammageing related mediators that were assessed in plasma using OLINK. Shown are boxplots with Normalized Protein Expression (NPX) values on a log2 scale. **c and d**) Multivariable linear regression models assessing the contribution of frailty, age, and sex to inflammageing-related cytokine levels (**c**) and tissue-resident memory T cells (**d**). Older fit and frail adults were included in a series of cytokine-specific regression models. Cytokine concentrations (IL-6, IL-8, HGF, TNF, CXCL9, and CCL3) were standardized (z-scored) prior to analysis. For each cytokine, a linear model was fitted with Frailty status (Frail vs Older Adult), age, and sex (M vs F) as predictors. Forest plots display the estimated beta coefficients with 95% confidence intervals for each predictor across all cytokines. Adjusted *p*-values (Benjamini–Hochberg correction) are shown beneath each estimate. Positive coefficients indicate higher cytokine levels associated with the predictor, whereas negative coefficients indicate lower levels. Panels are faceted by predictor term, illustrating how frailty, chronological age, and sex independently relate to inflammageing markers (**c**) or abundance of tissue-resident memory T cells (**d**) in the nasal mucosa. **e**) Spearman correlation matrix showing associations between tissue-resident memory T cell (TRM) populations and inflammageing-related soluble mediators detected in plasma from the same individuals. f) Spearman correlations between tissue resident memory T cell (Trm) populations and inflammageing related IL-6 detected in plasma from the same individuals. Individual points are shown. Statistical differences were assessed with Kruskal Wallis tests followed by Wilcoxon signed rank tests with Benjamini-Hochberg correction for multiple testing (A and B) \**P* < 0.05; ** *P < 0.01*, *** *P < 0.001*

**Supplementary Figure 5.**
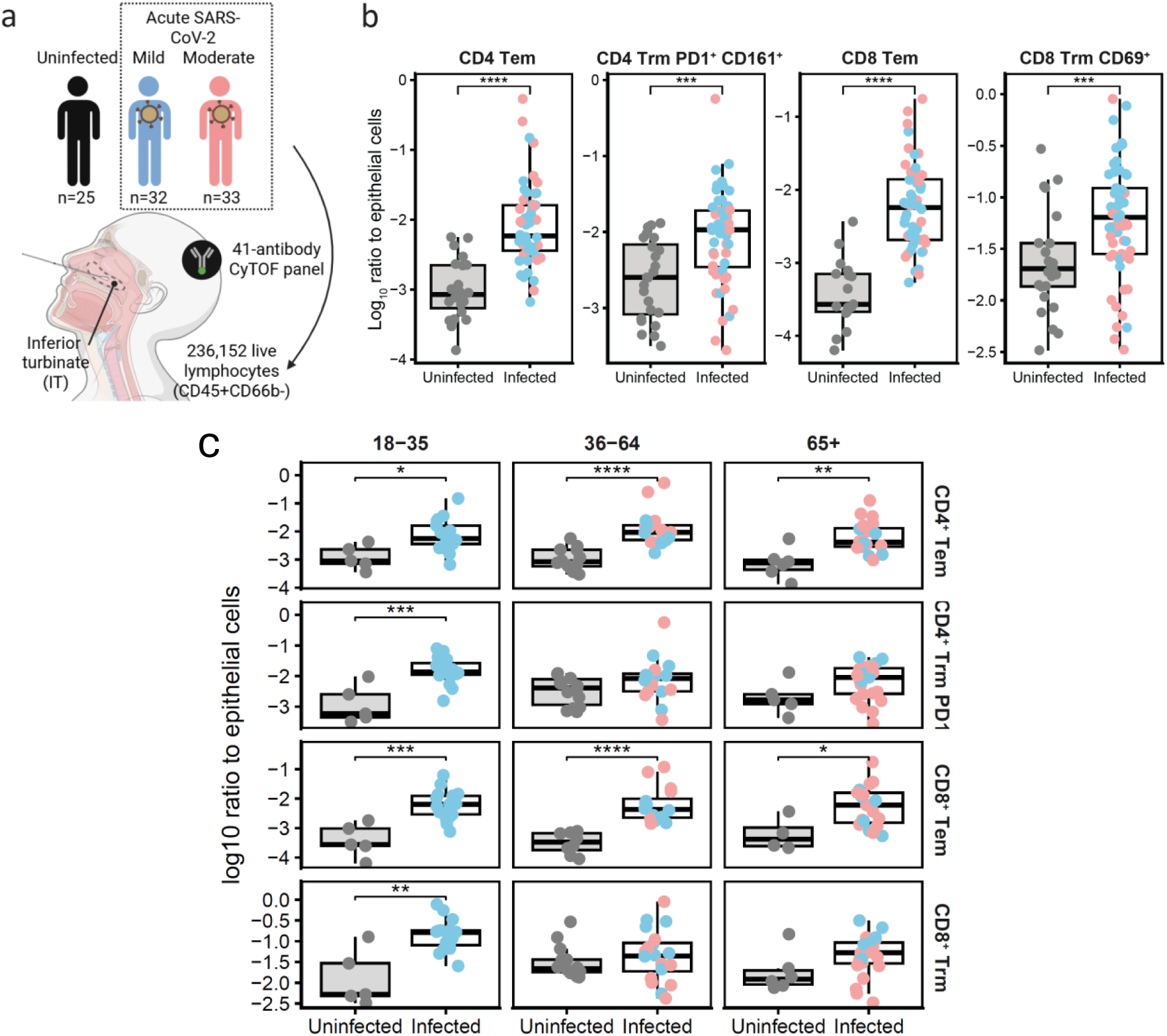
Age-related decline of immune cells in the upper-respiratory tract during acute SARS-CoV-2 infection. Nasal-derived cells were collected using curettage (RhinoPro©) and cryopreserved. **a**) Cross section of the upper respiratory tract, showing the anatomical site of sampling, the inferior turbinate. In total, 90 samples were thawed and incubated with a 41-antibody CyTOF panel and acquired on a Helios mass cytometer. **b**) T cell subsets were manually gated for CD4^+^ and CD8^+^ Tem (CD45RA^-^CCR7^-^) and Trm (CD69^+^) cells (Supplementary Table 4 and Extended Data Fig. 6). The most abundant CD4^+^ Trm was further defined based on expression of PD1 and CD161 to be in concordance with the CD103^+^CD4^+^ Trm population of Elbow cluster 10; Figure 1b. Ratio of these immune cell subsets normalized to the number of epithelial cells from the same sample are shown as boxplots and compared between uninfected and SARS-CoV-2 infected individuals. **c**) Boxplots are shown of the ratio of T cell subsets normalized to the number of epithelial cells from uninfected versus SARS-CoV-2 infected samples and stratified for age. Statistical differences were assessed with Mann-Whitney U tests (B and C) *P < 0.05; **P < 0.01; ***P < 0.001; ****P < 0.0001

**Supplementary Figure 6.**
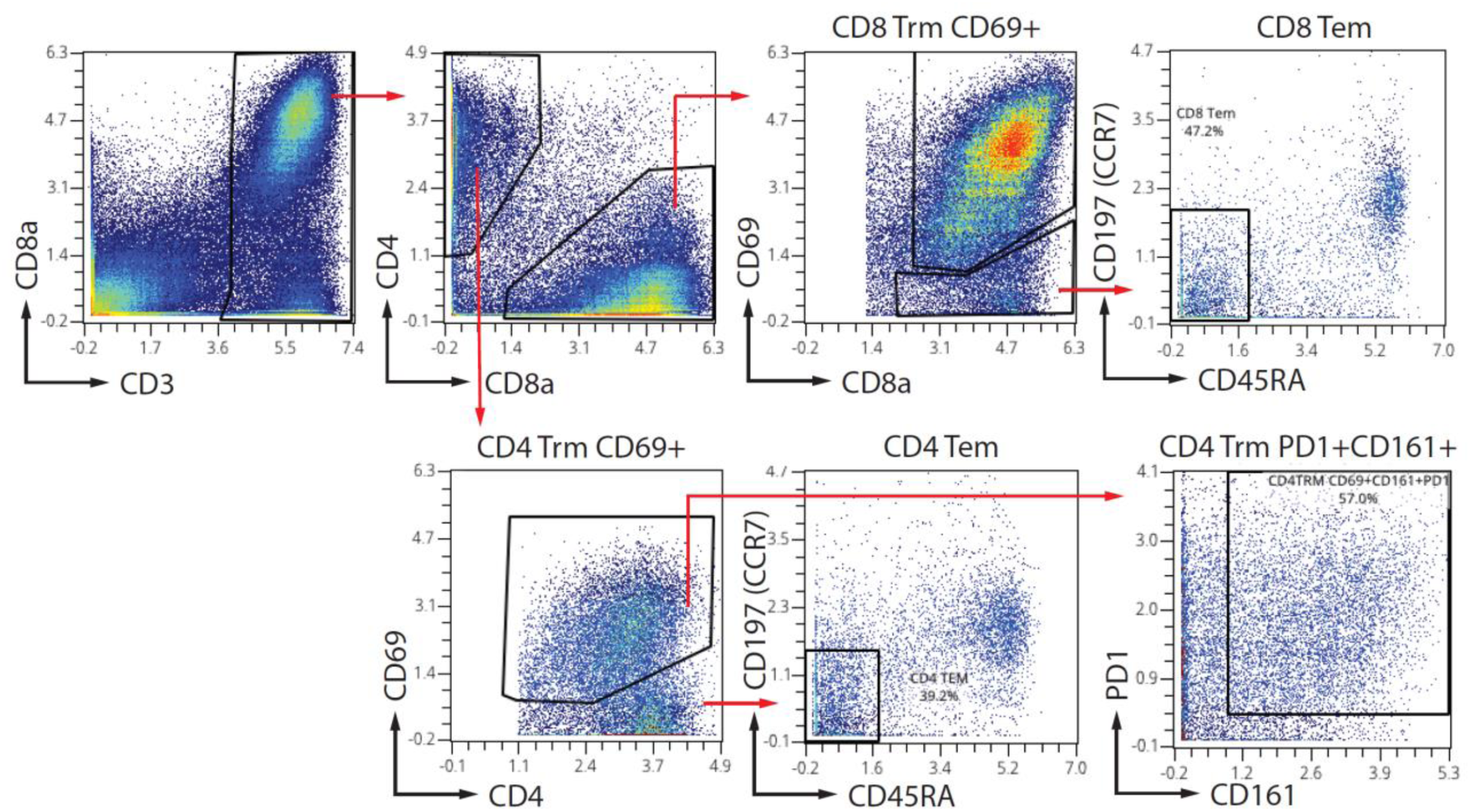
Gating strategy of tissue-resident T subsets from CyTOF acquired data. Nasal-derived samples were thawed, barcoded and measured in one single batch of twenty samples. Immune cells were manually gated and exported as .fcs files based on DNA dye and CD45 expression, with exclusion of cPARP positive apoptotic cells, as well as immune doublets (CD14+CD3+ , CD66b+CD3+ , CD14+CD66b+). Exported immune cell .fcs files were uploaded in OMIQ software for batch correction. Granulocytes (CD45+CD66b+) were gated out and remaining lymphocytes were further used for batch normalization. For batch normalization, lymphocytes were sub clustered using all markers except CD19/CD20, CD9 and EpCAM into 5 clusters using FLOWsom k=5 with a dimension of 15×15 and 25 training iterations, followed by Cytonorm with 101 quantiles to apply normalization across batches. T cell subsets were then manually gated in OMIQ.

**Supplementary Figure 7.**
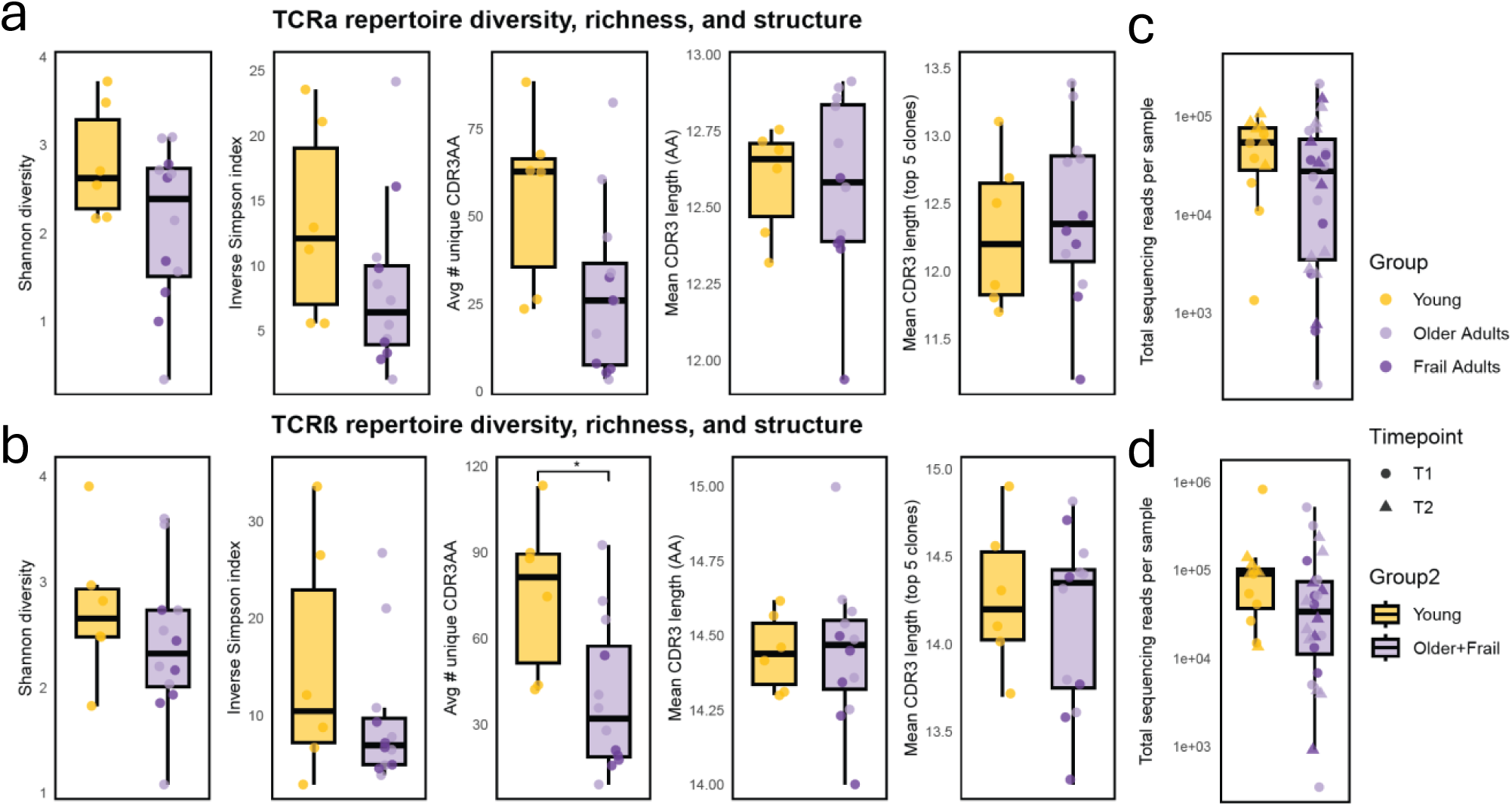
Age-Associated Differences in TCR Repertoire. Bulk TCR sequencing was performed to quantify clonal overlap and stability of T cells between paired samples collected at two timepoints. There were no respiratory infections reported between these timepoints. TCR repertoire characteristics were quantified after preprocessing, including removal of low-quality CDR3 sequences with non-productive rearrangements, extreme CDR3 lengths (>30 amino-acids) and low frequent clonotypes shared between all samples. Clonotype frequencies were normalized per sample, and only clonotypes contributing ≥0.01% of total reads were retained. **a and b**) For each participant, and TCRα (**a**) and TCRβ (**b**) separately, repertoire diversity (Shannon index, Inverse Simpson index), richness (Chao1), mean CDR3 amino-acid length (all clones and top 5 most abundant clones) and the number of unique CDR3 sequences. **c and d**) The total number of reads per sample and timepoint are shown for TCRα (**c**) and TCRβ (**d**), separately. Group comparisons were performed between young and older fit/frail adults using two-group Wilcoxon rank-sum tests, with Benjamini–Hochberg correction applied across metrics to control the false discovery rate. Adjusted p-values are shown on the plots.

**Supplementary Figure 8.**
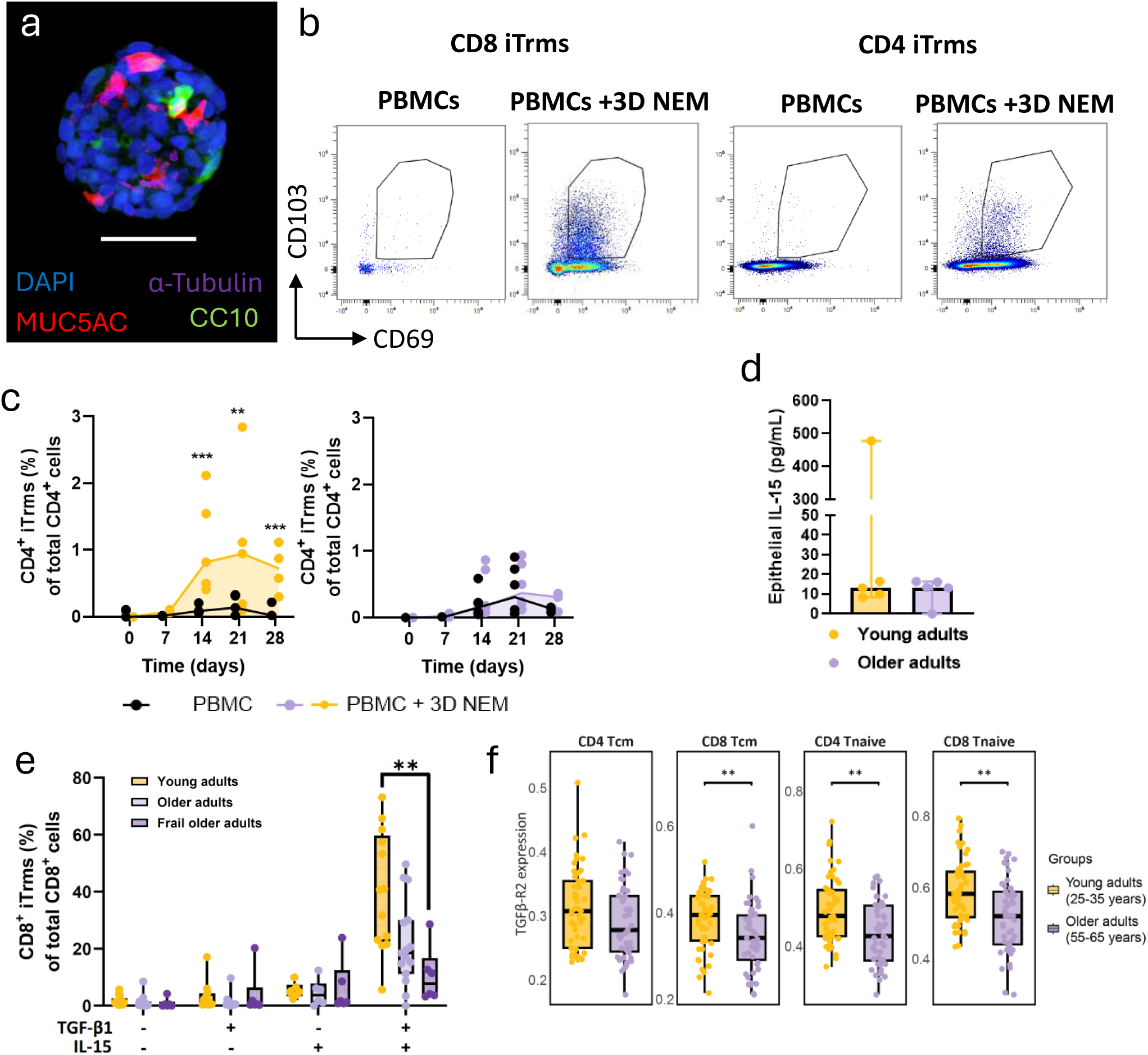
Impaired Tissue-resident memory T cell differentiation from older adults. The 3D nasal epithelial models (3D NEM) were generated from primary nasal epithelial cells collected from TINO participants using nasal curettage. After two weeks of differentiation, 3D NEMs were harvested and co-cultured in the presence or absence of freshly thawed autologous PBMCs. **a**) Confocal microscopy images of fixed and permeabilized 2 week old 3D nasal epithelial structure. The 3D nasal epithelial structures were stained with DAPI (blue, cell nuclei) and Muc5AC (red, goblet cells), acetylated alpha tubulin (purple, cilia/ciliated cells) and CC10 (green, club cells) fluorescently labelled antibodies. Z-stack imaging was used to capture the entire 3D structure. The 40x oil objective was used and the white scale bar indicates 40 μM. **b**) Representative FACS plots at day 14 showing the percentage of induced CD4^+^ and CD8^+^ tissue-resident memory T cells (iTrms) in presence and absence of 3D NEM. Cells were gated on live CD3⁺ T cells and analyzed for expression of tissue-resident markers (CD69 and CD103). **c**) Percentage of CD4^+^ iTrms over time are shown for PBMCs derived from young (n=5, yellow) and older adults (n=6, purple). **d)** Bar plots showing the production of IL-15 (pg/ml) of 3D NEMs from young (n=5) and older adults (n=5) at day 14. **e**) Barplots showing percentages of iTrms when PBMCs were cultured without 3D NEMs and supplemented with 5ng/ml TGF-β1 and/or 10ng/ml IL-15 **f)** Gene expression of the TGF-β receptor was reanalyzed from the publicly available SoundLife single-cell RNA-sequencing dataset. Four peripheral blood T-cell subsets were extracted based on the original cell annotations, and normalized expression values were visualized per sample. In total, 45 older adults (55-65 years) were compared to 49 young adults (25-35 years). Statistical differences were assessed with a linear mixed model ((log10iTRM%) ∼ day × condition) (C), Mann-Whitney U tests (D) or Kruskal Wallis tests followed by Wilcoxon signed rank tests with Benjamini-Hochberg correction for multiple testing (E and F) . *P < 0.05, **P < 0.01

**Supplementary Figure 9.**
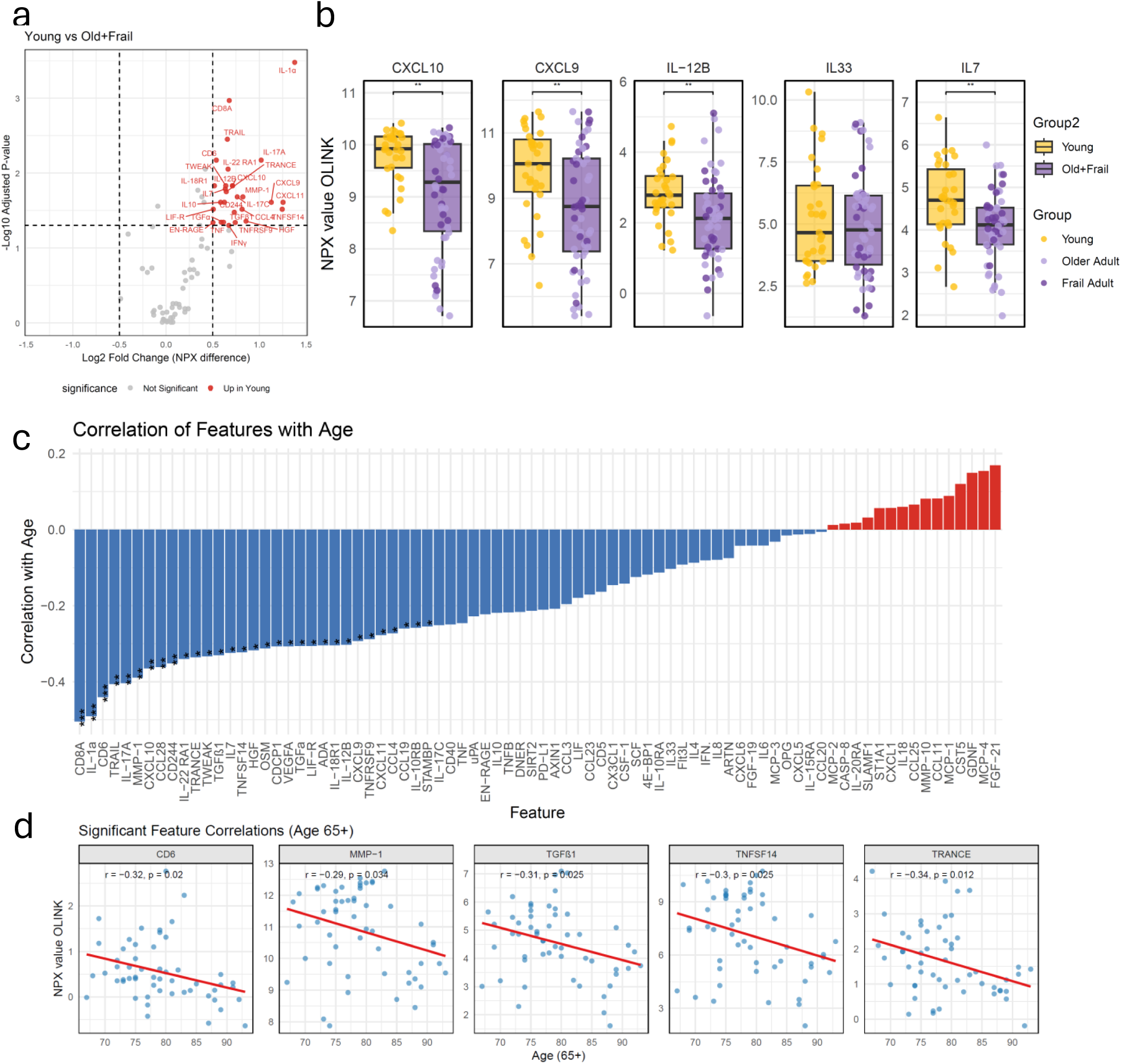
Inflammatory protein profiling using the Olink Target 96 Inflammation panel on nasal lining fluid. **a**) Volcano plots showing differential protein abundance between the indicated groups measured using the Olink Target 96 Inflammation panel. Protein levels are reported as normalized protein expression (NPX) values on a log2 scale. Each dot represents one of the 80 inflammatory proteins that passed quality control. **b**) Relative abundance of proteins associated with tissue-resident immune cell maintenance and induction. Protein levels are shown as NPX values (log2 scale). **c**) Spearman correlation analysis of inflammatory proteins with age across all study participants, including young, older, and frail individuals. Each bar indicates the correlation coefficient and asterixis indicate significant correlations after Benjamini–Hochberg correction for multiple testing. **d**) Shown are spearman correlation plots of inflammatory proteins that remained significant when analyzing participants aged ≥65 years only.

**Supplementary Figure 10.**
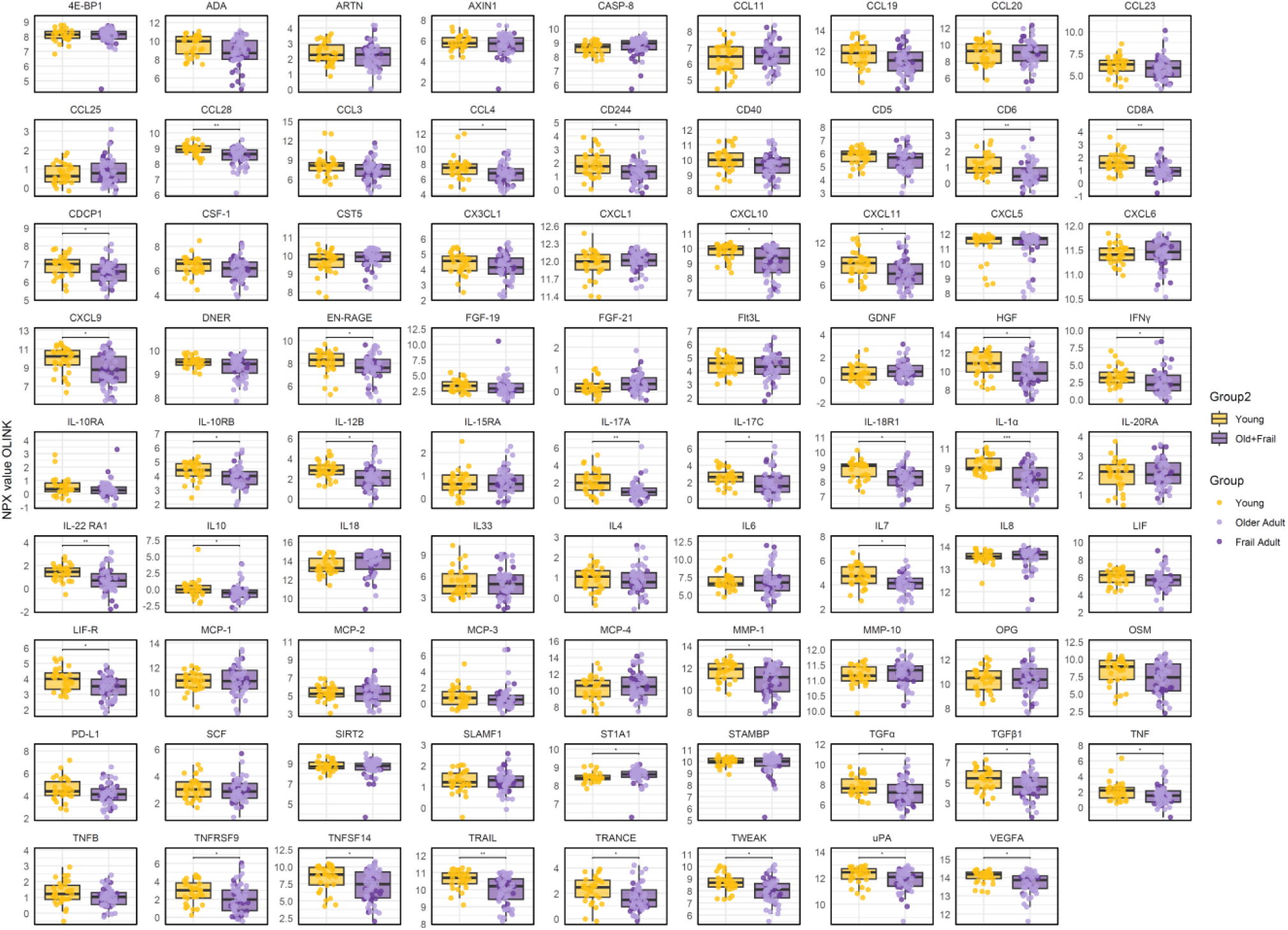
Inflammatory protein profiling using the Olink Target 96 Inflammation panel on nasal lining fluid. Distribution of 80 inflammatory proteins measured in the Olink Target 96 Inflammation panel shown as boxplots. Sixteen proteins did not meet quality control and were not further analyzed. Protein abundance is displayed as NPX values (log2 scale). Boxes represent the interquartile range with the median indicated, and whiskers indicate the 95%CI of the data.

**Supplementary Figure 11.**
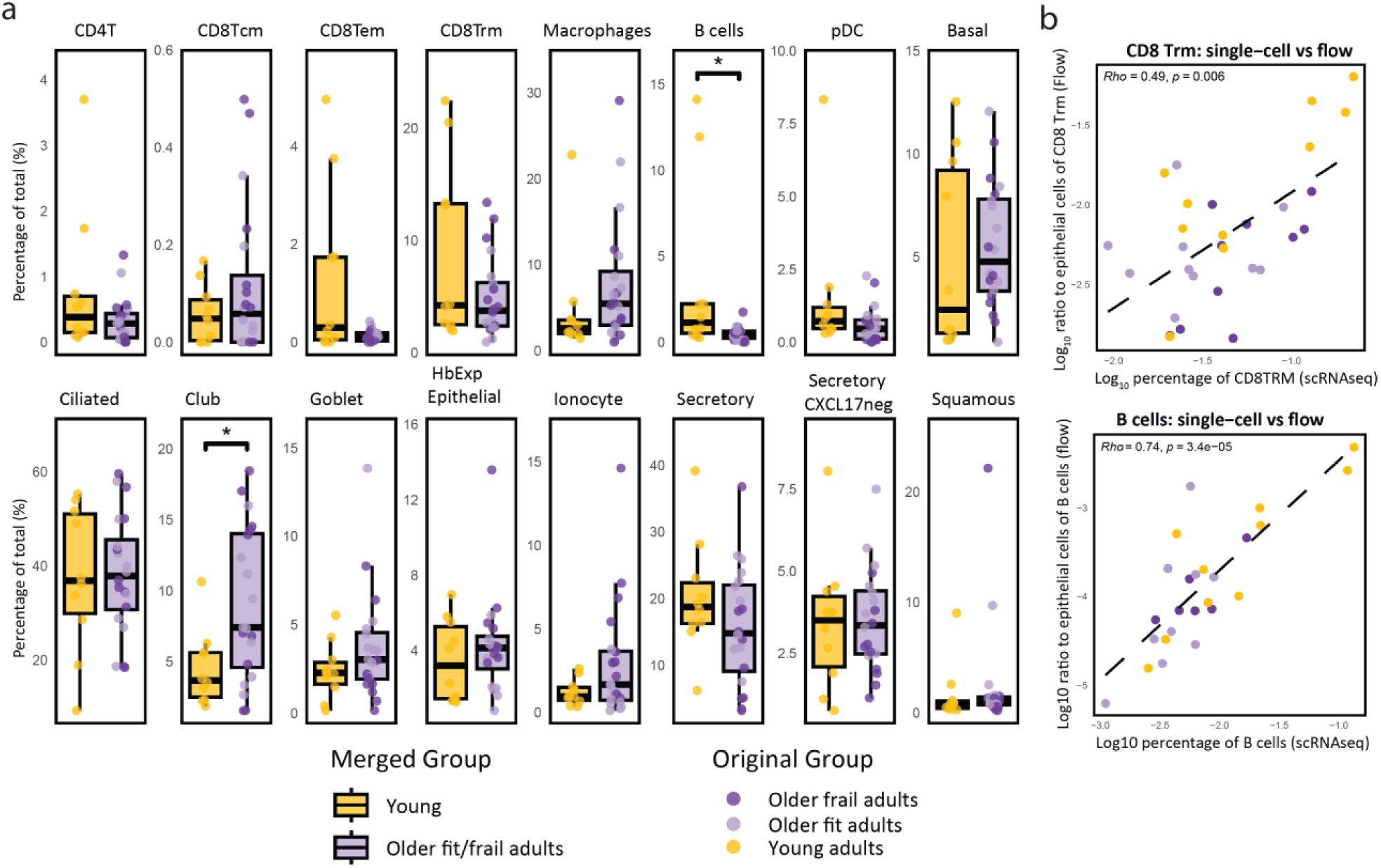
**Frequencies of nasal cell populations in young and older (fit/frail) adults based on single RNA sequencing**. **a**) Abundance of immune cell subsets and epithelial subsets expressed as a percentage of total cells. Data are shown as boxplots comparing young versus older fit/frail adults. Statistical differences between groups were assessed using the Wilcoxon rank-sum test. **b**) Scatter plots show the association between abundances of CD8 tissue-resident memory T (TRM) cells and B cell subsets quantified by single-cell RNA sequencing and spectral flow cytometry across samples. Each point represents an individual subject and is colored by group. Correlations were assessed using Spearman’s rank correlation coefficient, with the corresponding rho and p-values indicated in each panel. A linear regression line (dashed) is shown for visual guidance. Samples with zero values were excluded from the analysis.

**Supplementary Figure 12.**
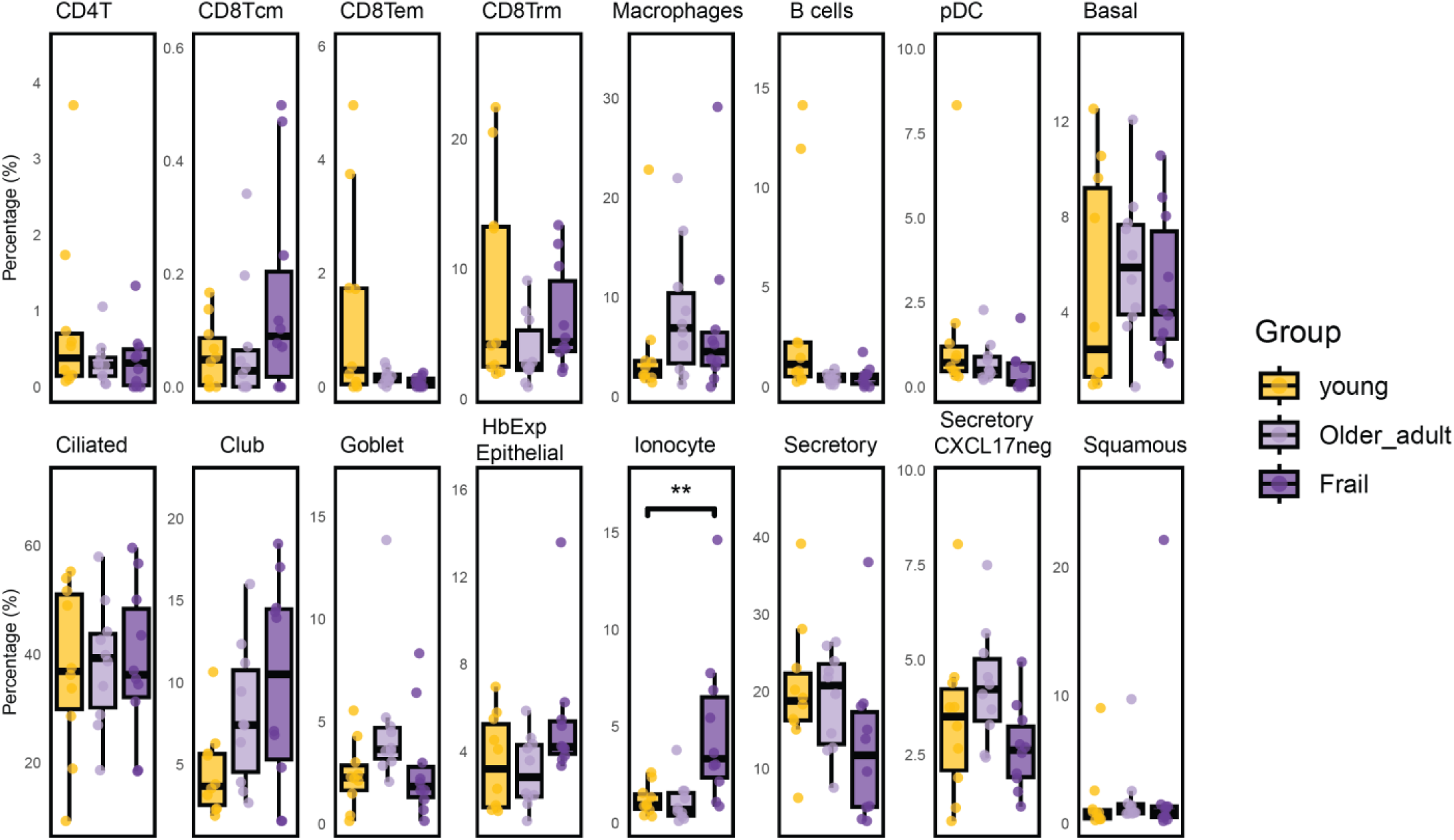
Frequencies of nasal cell populations in young, older fit, and older frail adults based on single-cell RNA sequencing. Top panel: relative abundance of immune cell subsets expressed as a percentage of total immune cells. Bottom panel: relative abundance of epithelial cell subsets expressed as a percentage of total epithelial cells. Data are shown as boxplots comparing the three age/fitness groups. Statistical differences were assessed using the Wilcoxon rank-sum test, and resulting p-values were adjusted for multiple testing using the Benjamini–Hochberg (BH) method.

**Supplementary Figure 13.**
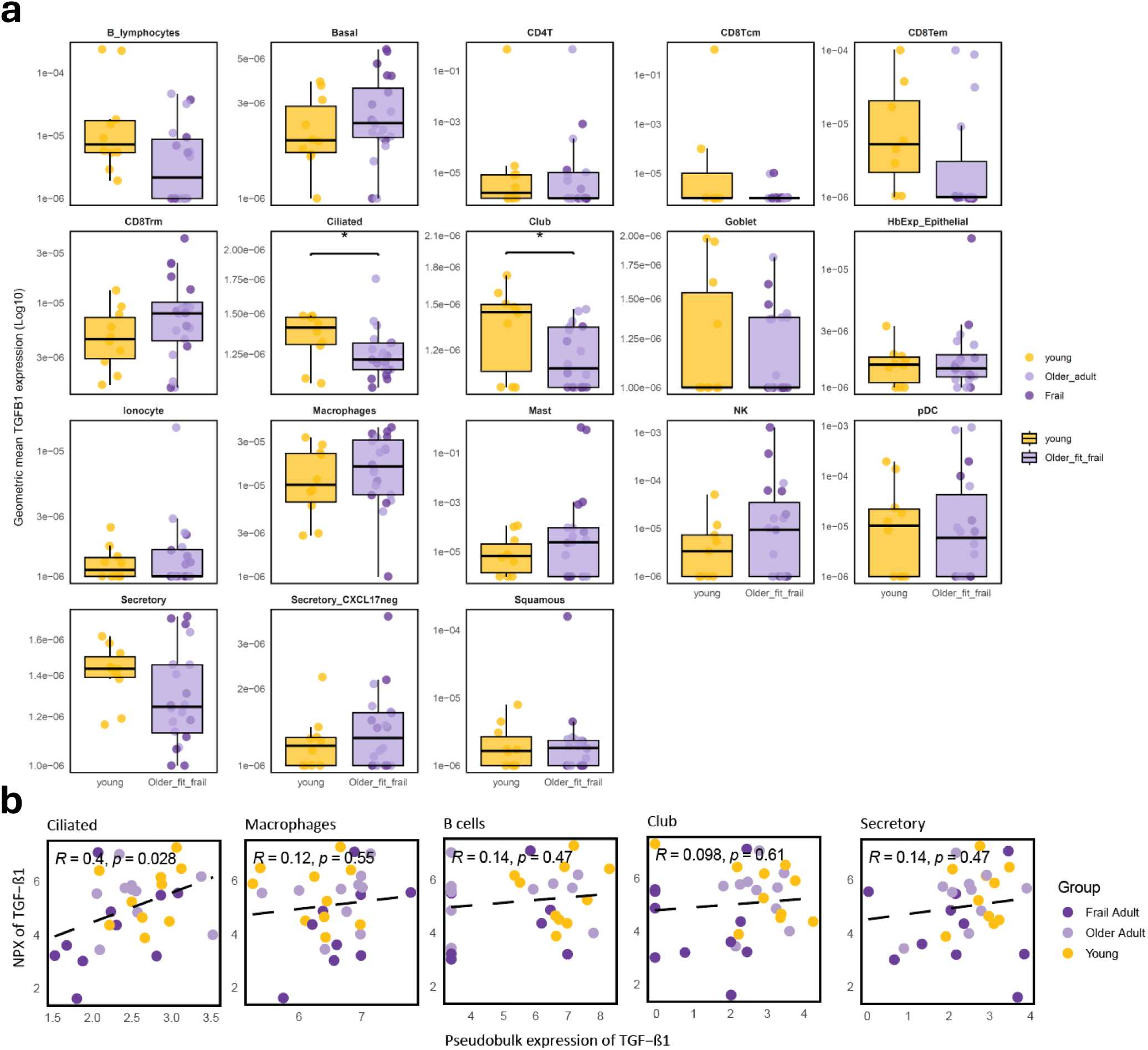
Expression of TGFB1⁺ across epithelial and immune subsets in the single-cell RNA-seq dataset. **a)**For each sample and cell type, TGFB1 expression was aggregated using the geometric mean, calculated as exp(mean(log(TGFB1_expr + 1e−6))). A small pseudocount (1e−6) was added to avoid log-transformation of zero values. Geometric means are plotted on a log10 scale to allow comparison across cell types with different expression ranges. Comparisons between young and older fit/frail individuals were performed using a Wilcoxon rank-sum test, applied per cell type. **b**) Scatterplots show the relationship between pseudobulk TGFB1 expression derived from single-cell RNA sequencing and nasal TGF-β1 protein levels measured using Olink (NPX) across five cell populations. Each point represents an individual sample, colored by group. Dashed lines indicate linear regression fits, and Pearson correlation coefficients (R) with corresponding p-values are shown in each panel. For the macrophage dataset, one outlier sample was excluded from the analysis to avoid disproportionate influence on the correlation.

**Supplementary Figure 14.**
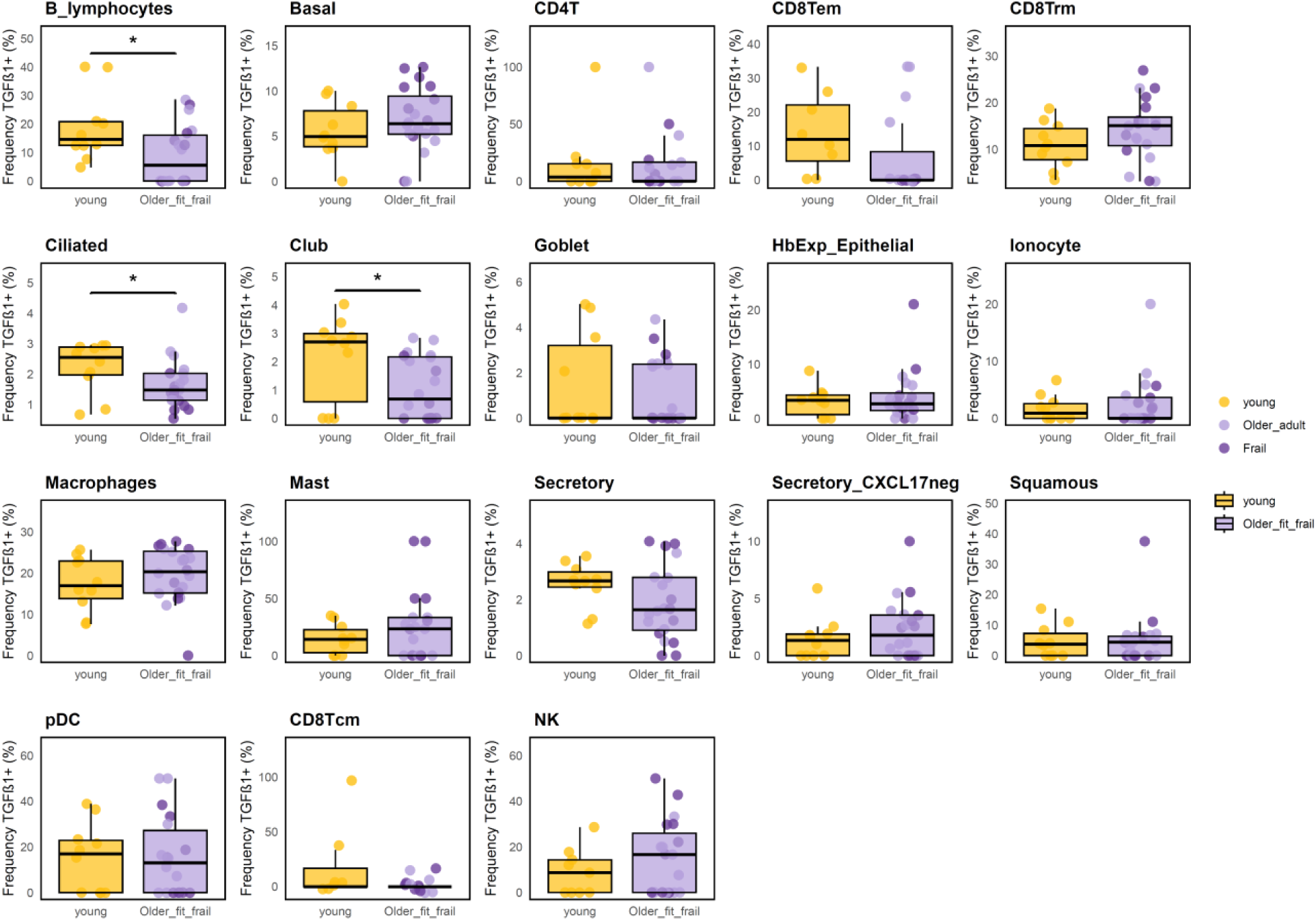
Frequency of TGFB1⁺ cells across epithelial and immune subsets in the single-cell RNA-seq dataset. For each sample and cell type, the proportion of TGFB1⁺ cells was quantified and visualized using boxplots. Comparisons between young and older fit/frail individuals were performed using a Wilcoxon rank-sum test, applied per cell type.

**Supplementary Figure 15.**
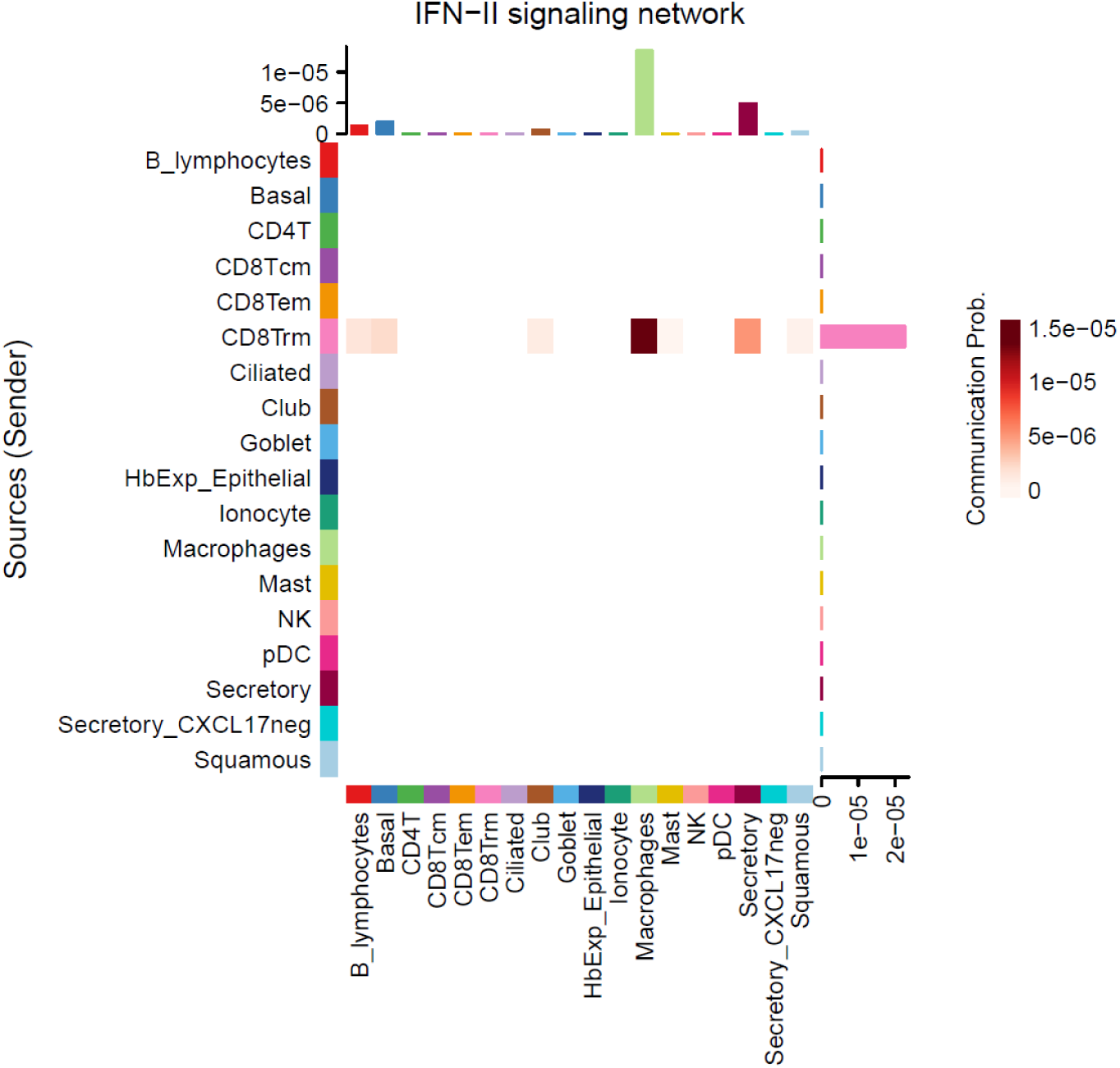
Comparison of intercellular signaling strength between groups using CellChat. Heatmap visualization of IFN-II (IFNG) signaling in nasal samples, generated using CellChat. The heatmap displays the contribution of individual ligand–receptor pairs to the IFN-II signaling pathway across cell types, with signal intensity shown in a red color scale. Pathway contribution analysis (bottom) quantifies the relative influence of each ligand–receptor interaction within the IFN-II network, highlighting the dominant signaling components driving this pathway in the dataset.

**Supplementary Figure 16.**
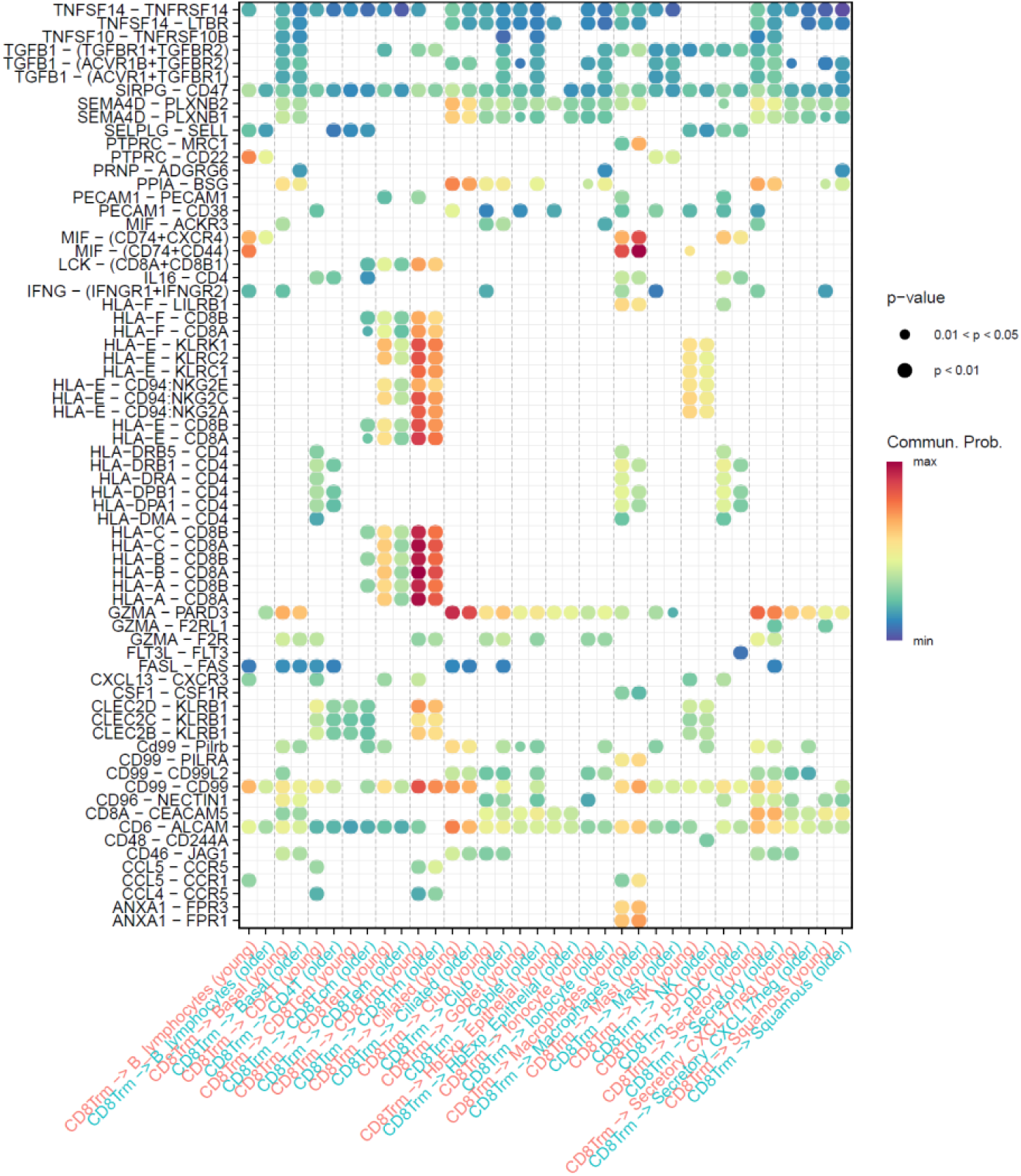
CellChat bubble plot showing ligand–receptor signaling involving CD8⁺ tissue-resident memory T cells (Trm). Outgoing and incoming interactions for CD8^+^ Trm are visualized and compared between groups (young (blue) vs older adults (red)). Each bubble represents a ligand–receptor pair, with bubble size proportional to communication probability and color indicating differential signaling strength. Only significant interactions detected in the comparison analysis are shown.

**Supplementary Figure 17.**
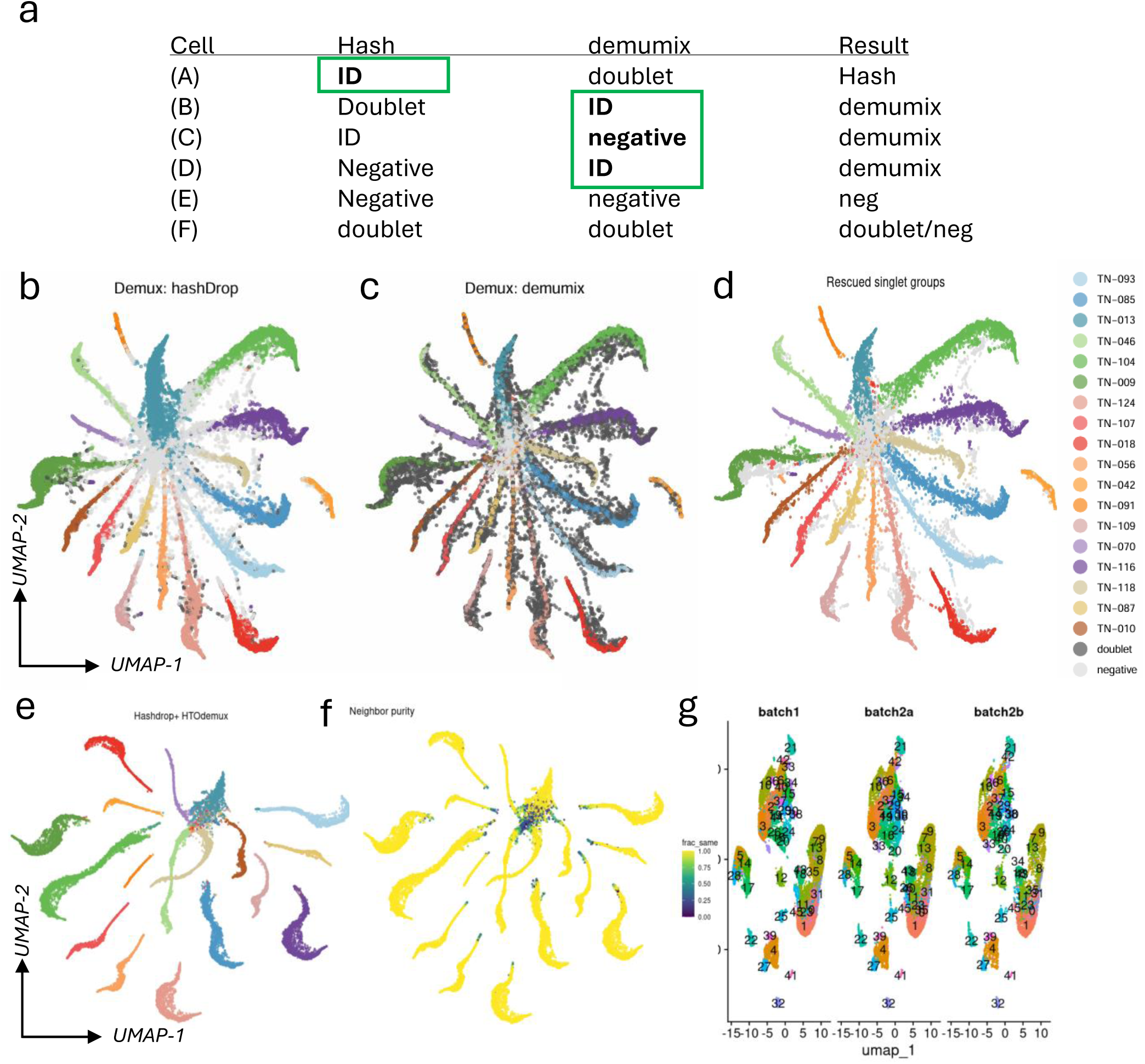
Donor demultiplexing, doublet removal, and neighborhood-purity filtering of HTO-labeled cells. **a**) Overview of demultiplexing strategy using HashDrop and HTODemux. Cells were assigned to donors using two independent demultiplexing approaches. **B and c**) Demultiplexing using hashDrop (B) and HTODemux (C). HTO count matrices were visualized using UMAP to assess sample identity and evaluate concordance between methods. **d**) Cells classified as doublets or negatives by both algorithms were intersected, and a consensus donor label was assigned to each cell. **e and f**) To further eliminate ambiguous donor assignments, we applied a k-nearest neighbor (kNN) neighborhood-purity filter in HTO count space using the RANN package (v2.6.2). For each cell, the proportion of its nearest neighbors sharing the same donor label was computed, and cells with neighborhood purity < 0.5 were excluded. **g**) Clusters were identified using the Harmony-corrected embeddings and visualized across batches to assess integration quality.

